# Altered activity of mPFC pyramidal neurons and parvalbumin-expressing interneurons during social interactions in a *Mecp2* mouse model for Rett syndrome

**DOI:** 10.1101/2024.08.06.606882

**Authors:** Destynie Medeiros, Likhitha Polepalli, Wei Li, Lucas Pozzo-Miller

## Abstract

Social memory impairments in *Mecp2* knockout (KO) mice result from altered neuronal activity in the monosynaptic projection from the ventral hippocampus (vHIP) to the medial prefrontal cortex (mPFC). The hippocampal network is hyperactive in this model for Rett syndrome, and such atypically heightened neuronal activity propagates to the mPFC through this monosynaptic projection, resulting in altered mPFC network activity and social memory deficits. However, the underlying mechanism of cellular dysfunction within this projection between vHIP pyramidal neurons (PYR) and mPFC PYRs and parvalbumin interneurons (PV-IN) resulting in social memory impairments in *Mecp2* KO mice has yet to be elucidated. We confirmed *social memory* (but not *sociability*) deficits in *Mecp2* KO mice using a new 4-chamber social memory arena, designed to minimize the impact of the tethering to optical fibers required for simultaneous *in vivo* fiber photometry of Ca^2+^-sensor signals during social interactions. mPFC PYRs of wildtype (WT) mice showed increases in Ca^2+^ signal amplitude during explorations of a novel toy mouse and interactions with both familiar and novel mice, while PYRs of *Mecp2* KO mice showed smaller Ca^2+^ signals during interactions only with live mice. On the other hand, mPFC PV-INs of *Mecp2* KO mice showed larger Ca^2+^ signals during interactions with a familiar cage-mate compared to those signals in PYRs, a difference absent in the WT mice. These observations suggest atypically heightened inhibition and impaired excitation in the mPFC network of *Mecp2* KO mice during social interactions, potentially driving their deficit in social memory.

## INTRODUCTION

Atypical social behaviors are characteristic of many neuropsychiatric disorders, including schizophrenia and neurodevelopmental disorders with autistic features (Brumback et al., 2018; Horiai et al., 2020; O’Tuathaigh et al., 2007; Piskorowski et al., 2016; Tao et al., 2022). Various brain regions have been associated with social behaviors (Bicks et al., 2020; Chen and Hong, 2018; Gunaydin et al., 2014; Ko, 2017; Montagrin et al., 2018), including the preference to interact with novel conspecifics over inanimate objects (i.e. sociability, social preference) and the memory of those conspecifics (i.e. social memory) (Chiang et al., 2018; Okuyama, 2018). However, the specific neuronal subtypes involved and their modulation remain to be fully uncovered. The hippocampus is well-known for its role in various types of memory (Benoy et al., 2018), including social memory. Pyramidal neurons (PYR) in area CA2 of the dorsal hippocampus (dHIP) and their projections to area CA1 of the ventral hippocampus (vHIP) are necessary for social memory (Hitti and Siegelbaum, 2014; Meira et al., 2018), while ventral CA3 PYRs are also implicated in social memory (Chiang et al., 2018). In addition to these intra-hippocampal connections, PYRs from ventral CA1 and subiculum that project to the nucleus accumbens (NAc) shell are also necessary for social memory (Okuyama et al., 2016). We recently demonstrated that either exciting or inhibiting PYRs in ventral CA1 that project to the medial prefrontal cortex (mPFC) using chemogenetics impairs social memory, without affecting sociability (Phillips et al., 2019).

Overall neuronal spiking activity within the mPFC network increases during social approach behaviors (Lee et al., 2016), and specific mPFC PYR ensembles that either increase or decrease their spiking activity during social interactions remain stable over repeated testing sessions (Liang et al., 2018). Moreover, a proper balance of excitatory and inhibitory synaptic inputs onto mPFC excitatory PYRs is crucial for sociability: selective optogenetic manipulation of either PYRs or their inhibitory inputs from local GABAergic interneurons (INs) results in reversible sociability deficits in mice (Yizhar et al., 2011). Likewise, optogenetic stimulation of parvalbumin (PV) expressing INs in the mPFC can partially rescue the social memory deficits caused by optogenetic inhibition of vHIP neurons (Sun et al., 2020). Additionally, either optogenetic or chemogenetic stimulation of mPFC PV-INs rapidly triggers social approaches, promoting sociability, as well as rescuing impaired sociability caused by social isolation during adolescence (Bicks et al., 2020). These findings suggest a critical role of proper mPFC network activity in sociability and social memory, specifically when modulated by PV-INs.

Among monogenetic neurodevelopmental disorders with autistic features, Rett syndrome (RTT) is a debilitating condition that presents with a multitude of neurological and psychiatric signs and symptoms, including motor control deficits, breathing irregularities, anxiety, panic attacks, seizures, autistic features, speech problems, and intellectual disability (Chahrour and Zoghbi, 2007; Haas, 1988; Le Bras, 2022). RTT primarily affects females, occurring in approximately 1 in 10,000 births (Armstrong, 2005; Laurvick et al., 2006). Initially, individuals with RTT develop typically until 6-18 months of age, when they begin to regress in cognitive skills and language, accompanied by the onset of multiple neuropsychiatric symptoms (Glaze et al., 2010; Hagberg et al., 1983). The majority of classic RTT cases are caused by mutations in the X-linked gene *MECP2* (Amir et al., 1999). Methyl-CpG binding protein 2 (MeCP2) is a nuclear protein that binds to methylated DNA, functioning as a transcriptional modulator (Tillotson and Bird, 2020). Mice deficient in *Mecp2* exhibit multiple neurological phenotypes that are analogous to signs and symptoms observed in RTT individuals (Chen et al., 2001).

Of relevance to seizure disorders and intellectual disability in RTT individuals, disinhibition of principal excitatory neurons in hippocampal area CA3 causes hippocampal network hyperactivity in *Mecp2* knockout (KO) mice (Calfa et al., 2011; Calfa et al., 2015), which leads to enhanced excitatory synaptic transmission that occludes long-term potentiation at CA3-CA1 synapses (Li et al., 2016), likely contributing to the seizure phenotype in mice (D’Cruz et al., 2010; Wither et al., 2018; Zhang et al., 2008) and RTT individuals (Glaze et al., 2010). Similarly, brainstem nuclei responsible for breathing control also show hyperactivity due to atypically heightened excitatory synaptic transmission (Dhingra et al., 2013; Katz et al., 2009). Conversely, cortical networks show hypoactivity due to either enhanced inhibition or reduced excitation (Dani et al., 2005; Durand et al., 2012), including the mPFC (Phillips et al., 2019; Sceniak et al., 2016). Despite such local hypoactivity in the mPFC, stimulation of afferent inputs from the vHIP evokes larger, longer, and more widespread depolarizations in the mPFC of *Mecp2* KO mice, which is accompanied by social memory deficits, without affecting sociability (Phillips et al., 2019). Furthermore, chemogenetic inhibition of mPFC-projecting vHIP PYRs for 3 weeks normalized vHIP-driven mPFC activity and improved social memory (Phillips et al., 2019), suggesting that heightened hippocampal activity due to CA3 disinhibition spreads to the mPFC via this monosynaptic excitatory long-range projection. These vHIP afferent axons synapse on mPFC PYRs (Dembrow et al., 2015; Little and Carter, 2012; Liu and Carter, 2018; Phillips et al., 2019), as well as onto different subclasses of GABAergic INs, including those expressing PV, somatostatin (SOM), calretinin (CR), and cholecystokinin (CCK) (Liu and Carter, 2018; Liu et al., 2020b; Marek et al., 2018; Phillips et al., 2019), with the fraction of vHIP axons synapsing onto PV-INs and pyramidal tract PYRs showing the most significant differences between *Mecp2* KO mice and their littermate wildtype (WT) controls (Phillips et al., 2019). Consistent with the role and engagement of the mPFC during social behaviors (Lee et al., 2016; Liang et al., 2018; Yizhar et al., 2011), *Mecp2* KO mice show deficits in social memory that are rescued by chemogenetic inhibition of mPFC-projecting ventral CA1 PYRs (Phillips et al., 2019).

Here, we used fiber photometry of intracellular Ca^2+^ signals as surrogates of neuronal spiking activity, and show that mPFC PYRs of WT mice have increased amplitude of activity at the onset of explorations of a toy mouse, and interactions with both familiar and novel mice, while PYRs of *Mecp2* KO mice showed smaller Ca^2+^ signals only during interactions with live mice. On the other hand, mPFC PV-INs of *Mecp2* KO mice showed larger Ca^2+^ signals during interactions with a familiar cage-mate compared to those signals in PYRs, a difference absent in WT mice. These observations indicate atypically heightened inhibition from a specific subclass of PYR-targeting GABAergic IN (i.e. PV-INs), as well as impaired excitatory PYR activity in the mPFC of *Mecp2* KO mice during social interactions, potentially driving their deficit in social memory. Understanding the dynamic interplay between PYRs and different classes of GABAergic INs in the mPFC that are targets of vHIP inputs during social interactions is crucial for developing rational therapies for RTT and other neurodevelopmental disorders with social memory deficits, including those with autistic features.

## MATERIAL AND METHODS

### Mice

Breeding pairs of mice lacking exon 3 of the X chromosome-linked *Mecp2* gene (B6.Cg-*Mecp2*^tm1.1Jae^, “Jaenisch” strain in C57BL/6 background; stock number: 000415-UCD) (Chen et al., 2001) were purchased from the *Mutant Mouse Regional Resource Center* at the University of California, Davis. A colony was established at the University of Alabama at Birmingham (UAB) by mating WT males with heterozygous *Mecp2*^tm1.1Jae^ females, as recommended by the supplier; genotyping was performed by PCR of DNA samples from tail clips. Mice for all experiments were male hemizygous *Mecp2*^tm1.1Jae^ (i.e. *Mecp2* KO), and develop typically until 5–6 weeks of age (P35-P42), when they begin to exhibit RTT-like motor symptoms, such as hypoactivity, hind limb clasping, and reflex impairments. PV-Cre mice were purchased from The Jackson Laboratory (strain #008069); male *Mecp2* KO::PV-Cre mice and their male WT::PV-Cre littermates for experiments were obtained by mating female heterozygous *Mecp2*^tm1.1Jae^ mice with male PV-Cre mice. Animals were handled and housed according to the *Committee on Laboratory Animal Resources* of the National Institutes of Health. All experimental protocols were annually reviewed and approved by the *Institutional Animals Care and Use Committee* of UAB. One week before and throughout the experiments, mice were kept in an IACUC-approved dedicated room inside our lab (to avoid transportation stress) with a 12h light:12h dark cycle with lights on at 6am and off at 6pm.

### Intracranial virus injections and fiber optic implantation

Mice were anesthetized with 4% isoflurane vapor in 100% oxygen, and maintained with 1-2% isoflurane vapor in 100% oxygen. Mice were fixed in a stereotaxic frame (Kopf Instruments, Tujunga, CA), and their body temperature was maintained with a heating pad. For fiber photometry of Ca^2+^ sensors, 300nL of AAV1-CamKIIa-jGCaMP8m-WPRE (Addgene, #176751) to target PYRs, AAV1-Syn-FLEX-jGCaMP8f-WPRE (Addgene, #162379) to target PV-INs, or AAV1-Syn-FLEX-jRGECO1a-WPRE (Addgene #100853) were injected unilaterally into the prelimbic region of the mPFC of male *Mecp2* KO::PV-Cre mice and their male WT::PV-Cre littermates at postnatal day (P)-23-25 (1.45 A/P, 0.5 M/L, and -1.45 D/V from bregma) at a rate of 20nL/min using a microsyringe pump (UMP3 UltraMicroPump, Micro4; World Precision Instruments, Sarasota, FL). A 1.25mm ceramic ferrule (core 200µm, NA 0.39, length 2mm, RWD) was later implanted at the same stereotaxic coordinates, and fixed to the skull with quick adhesive cement (C&B-Metabond), and the exposed skull was covered with dental cement.

### Social behaviors in new 4-chamber social arena

Our 4-chamber social arena was adapted from a commercial *SocioBox* apparatus, which has 1 central chamber separated by transparent perforated partitions from 5 side chambers (Krueger-Burg et al., 2016). We redesigned it by removing 2 of the side chambers to simplify choices for the test mouse placed in the center chamber. Mice freely navigate the central chamber while different stimuli are placed in the 3 side chambers, which included an inanimate social object (toy mouse), a sex- and age-matched cage-mate mouse, and a sex-matched age-matched novel mouse. The partitions have 31 holes that allow for whisker and odor exchange between the experimental test mouse and 3 different stimuli. The experimental design involved a 2-day habituation period, during which mice were acclimated to the center chamber of the 4-chamber social arena for 5 min each day. Following habituation, mice underwent 3 consecutive trials. First, in the habituation phase (5 min) mice were allowed to explore the central chamber. Second, in the *sociability* phase (5 min) a social object and a sex-matched cage-mate were introduced into separate side chambers, and test mice were allowed to explore both social stimuli for 5 min to assess preference for a live mouse (i.e. sociability, social preference). Last, in the *social memory* phase (5 min) a novel sex- and age-matched mouse was introduced into the 3rd side chamber, and test mice were allowed to explore all 3 social stimuli for 5 min to assess preference for social novelty (i.e. social memory). All sessions were performed under minimal red light illumination from dim LED headbands, and recorded from above at 40 frames-per-second (fps) with an IR-sensitive GigE Camera (scA780-54gm, Basler) controlled by *Ethovision* XT17 (Noldus). We quantified the time spent within an interaction zone of 2.5cm from the perforated partition defined in *Ethovision*.

### Fiber photometry acquisition and data analysis

Fiber photometry recordings were performed by using a sCMOS-based system (FP3002, Neurophotometrics Ltd, San Diego, CA), which uses the open source software *Bonsai* (Lopes et al., 2015; Martianova et al., 2019). After allowing 23-25 days of recovery from virus injections and cannula implantation, P47-50 mice were habituated to the handling necessary for connecting the patch cord-fiber optic tethering, and allowed to explore the 4-chamber arena for 2 consecutive days; social behavior testing started the following day. jGCaMP8m was excited with 470nm (Ca^2+^ sensitive) and 415nm (Ca^2+^ insensitive, i.e. isosbestic) light from LEDs, and its fluorescence emission at 494-531nm was sampled at 40 fps with a sCMOS camera. Dual-color photometry from different neuronal populations in the same mouse was performed using jGCaMP8m as above, and jRGECO excited with 560nm light , with its emission detected at 586-627nm by the same sCMOS. Photometry signals were continuously recorded during the entire 5min of each phase of the 4-chamber social test. Transistor-transistor logic (TTL) signals were obtained from *Ethovision* to produce timestamps of the start and stop of each toy exploration or mouse interaction event in register with fiber photometry signals in *Bonsai*, as well as for the quantification of the time spent within the interaction zone of 2.5cm from the perforated partition. Each toy exploration or mouse interaction event started when the test mouse entered the interaction zone (2.5cm from the perforated partition facing each of the side chambers), and defined as time *zero*. To include mPFC Ca^2+^ sensor signals known that precede social interactions in similar assays (Bicks et al., 2020), we further divided that preceding time into a 2-second *Baseline* epoch (-5s to -3s) and a 2-second *Pre* exploration/interaction epoch (-2s to 0s); signals were also analyzed during a 2-second *Post* exploration/interaction epoch (0 to 2s).

Photometry traces during those events were analyzed using custom written code in *Python* (Martianova et al., 2019), based on a isosbestic method (Kim et al., 2016). First, the traces of the intensity of Ca^2+^ sensor emission when excited with 470nm (Ca^2+^ sensitive) and 415nm (isosbestic Ca^2+^ insensitive) during each event were smoothed with a *moving mean* algorithm. Second, the slow slope (sensor bleaching) and low frequency fluctuations (due to tissue motion and fiber bending) in those traces were removed from each emission channel with the *adaptive iteratively re-weighted Penalized Least Squares* (airPLS) algorithm (https://github.com/zmzhang/airPLS). Third, those baseline-corrected traces from each emission channel were standardized (‘z-scored’) using their mean and standard deviation. Fourth, the z-score of the Ca^2+^ insensitive signal (415nm) trace was fitted to the z-score of the Ca^2+^ sensitive signal (470nm) trace by non-negative linear regression. Finally, the z-score of the Ca^2+^ sensitive trace was divided by the regression-fitted z-score of the Ca^2+^ insensitive trace to obtain a single trace of the z-scored fractional change in Ca^2+^ sensitive emission intensity (deltaF_470_/F_410_), per mouse. We used the maximum value (peak) of the z-scored deltaF/F as a proxy of both the number of neurons spiking near the fiber and the intracellular Ca^2+^ levels reached by each of those neurons. We calculated the area-under-the-curve (AUC) of the z-scored deltaF/F trace using Simpson’s Rule, and used it to estimate the temporal structure (i.e. dynamics) of neuronal population spiking near the fiber (i.e. synchronization). At the conclusion of the experiments, all mice underwent transcardial perfusion with phosphate buffer saline (PBS, 100mM) followed by 4% paraformaldehyde (in 100mM PBS) to confirm cannula placement and expression of Ca^2+^ sensors. Mice with incorrect fiber placement and/or expression of Ca^2+^ sensors were excluded from further analyses.

### Statistical analyses

All experiments were conducted with at least 3 independent cohorts. Statistical analyses were performed using *Python* and *Prism* v10 (GraphPad). All data presented as mean +/- SEM. Statistical significance conventions are: *: p<0.05; **: p<0.01; ***: p<0.001.

## RESULTS

The initial validation of our new 4-chamber social arena was done using naive untethered male WT mice (Fig. 1). All other experiments (Figs. 2-6) were performed in *Mecp2* KO mice crossed with PV-Cre mice (referred to as *Mecp2* KO::PV-Cre) and their WT littermates (referred to as WT::PV-Cre), all of which expressed Ca^2+^ sensors in either mPFC PYRs or PV-INs, and had fiber-optic cannulas implanted in the mPFC for fiber photometry of intracellular Ca^2+^ sensors during behavioral tasks in the novel 4-chamber social arena.

**Figure 1:**
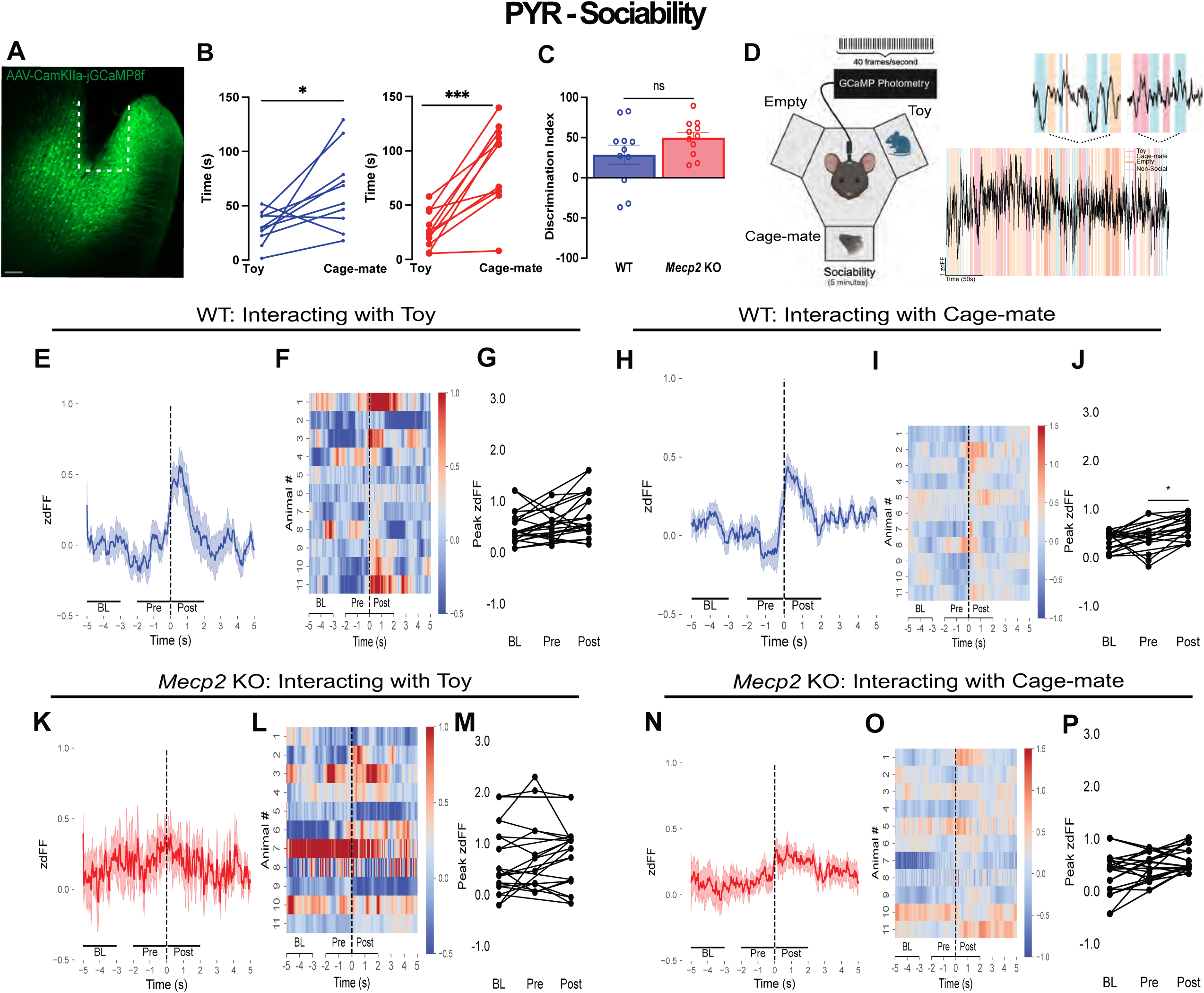
The spiking activity of mPFC PYRs from WT::PV-Cre mice respond to social stimuli, which is absent in *Mecp2* KO::PV-Cre mice. (A) Representative histological image of GCaMP8m and fiber placement (dotted line) in the mPFC. Scale bar, 100 µm. (B) Time spent interacting with the toy or cage-mate during the sociability trial of the 4-Chamber social interaction test for WT::PV-Cre(left) and *Mecp2* KO::PV-Cre mice (right) (C) Discrimination indices of sociability trial of WT::PV-Cre and *Mecp2* KO::PV-Cre mice (D) Schematic of sociability trial and a representative fiber photometry trace, zoomed in during interaction with the toy and the cage-mate (E) Calcium traces averaged across WT::PV-Cre animals during behavioral interactions with the toy mouse. Dark blue line denotes average signal, shaded blue denotes SEM. Time zero denotes behavioral bout onset (F) Heatmap depicting individual calcium response during behavioral interactions with the toy mouse of WT::PV-Cre mice (G) Peak mPFC PYR activity for each WT::PV-Cre mouse comparing Baseline (BL) (-5 to -3s before bout onset) Pre (-2 to 0s) and Post (0 to 2s) (H) Calcium traces averaged across WT::PV-Cre animals during behavioral interactions with the cage-mate. Dark blue line denotes average signal, shaded blue denotes SEM. Time zero denotes behavioral bout onset (I) Heatmap depicting individual calcium response during behavioral interactions with a littermate conspecific of WT::PV-Cre mice (J) Peak mPFC PYR activity for each WT::PV-Cre mouse comparing Baseline (BL) (-5 to -3s before bout onset) Pre (-2 to 0s) and Post (0 to 2s) (K) Calcium traces averaged across *Mecp2* KO::PV-Cre mice during behavioral interactions with the toy mouse. Dark red line denotes average signal, shaded red denotes SEM. Time zero denotes behavioral bout onset (L) Heatmap depicting individual calcium response during behavioral interactions with the toy mouse of *Mecp2* KO::PV-Cre mice (M) Peak mPFC PYR activity for each *Mecp2* KO::PV-Cre mouse comparing Baseline (BL) (-5 to -3s before bout onset) Pre (-2 to 0s) and Post (0 to 2s) (N) Calcium traces averaged across *Mecp2* KO::PV-Cre mice during behavioral interactions with the cage-mate. Dark red line denotes average signal, shaded red denotes SEM. Time zero denotes behavioral bout onset (O) Heatmap depicting individual calcium response during behavioral interactions with a cage-mate of *Mecp2* KO::PV-Cre mice (P) Peak mPFC PYR activity for each *Mecp2* KO::PV-Cre mouse comparing Baseline (BL) (-5 to -3s before bout onset) Pre (-2 to 0s) and Post (0 to 2s). Data are mean ±SEM.

**Figure 2:**
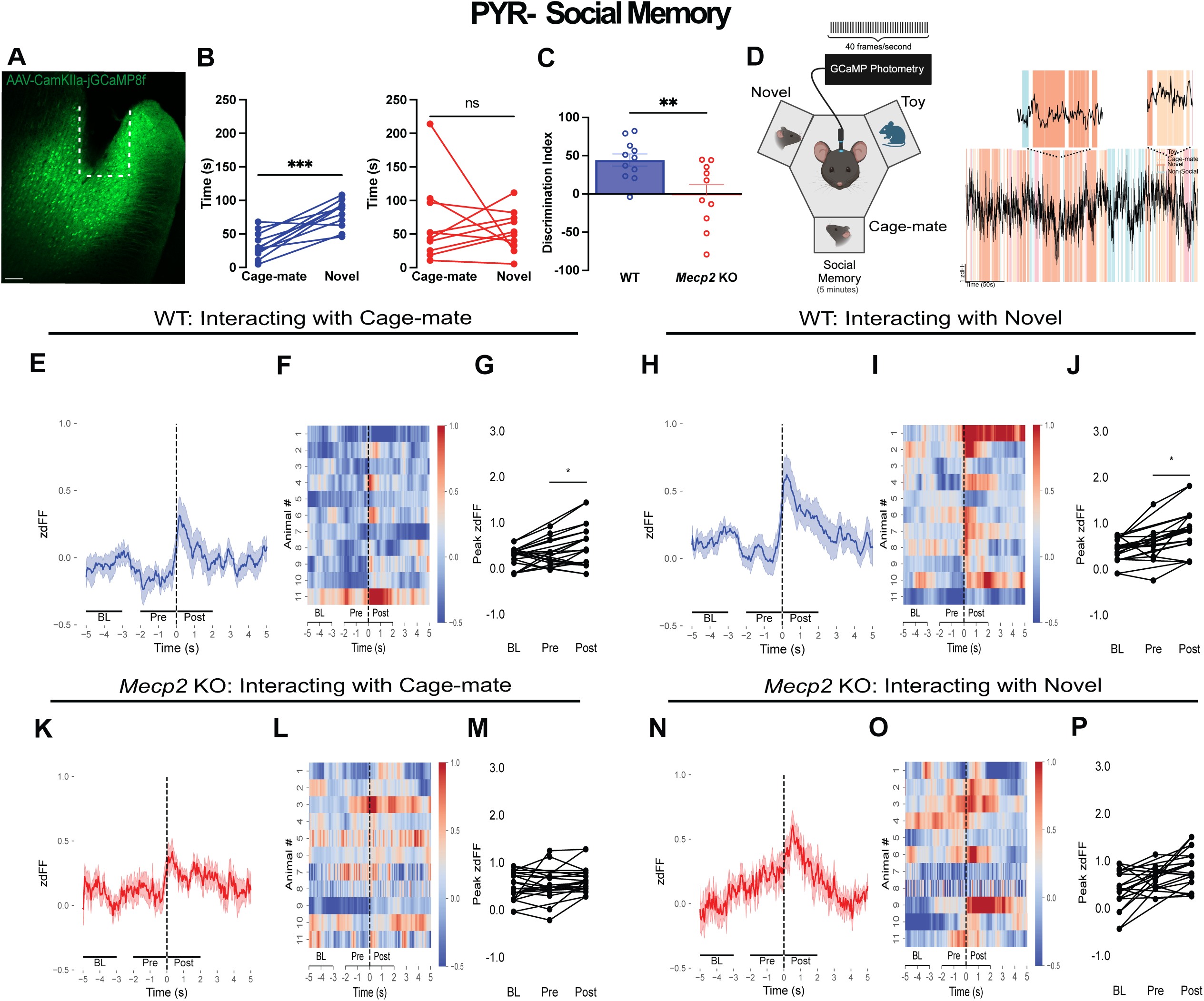
The spiking activity of mPFC PYRs during social interactions is lower and more temporally spread-out in *Mecp2* KO::PV-Cre mice than in WT::PV-Cre mice. (A) Representative histological image of GCaMP8m and fiber placement (dotted line) in the mPFC. Scale bar, 100 µm. (B) Time spent interacting with the cage-mate or novel mouse during the social memory trial of the 4-Chamber social interaction test for WT::PV-Cre (left) and *Mecp2* KO::PV-Cre mice (right) (C) Discrimination indices of the social memory trial of WT::PV-Cre and *Mecp2* KO::PV-Cre mice (D) Schematic of social memory trial and a representative fiber photometry trace, zoomed in during interaction with the cage-mate and the novel mouse (E) Calcium traces averaged across WT::PV-Cre animals during behavioral interactions with the cage-mate. Dark blue line denotes average signal, shaded blue denotes SEM. Time zero denotes behavioral bout onset (F) Heatmap depicting individual WT::PV-Cre mouse calcium response during behavioral interactions with the cage-mate (G) Peak mPFC PYR activity for each WT::PV-Cre mouse comparing Baseline (BL) (-5 to -3s before bout onset) Pre (-2 to 0s) and Post (0 to 2s) (H) Calcium traces averaged across WT::PV-Cre animals during behavioral interactions with a novel conspecific. Dark blue line denotes average signal, shaded blue denotes SEM. Time zero denotes behavioral bout onset (I) Heatmap depicting individual calcium response during behavioral interactions with a novel conspecific of WT::PV-Cre mice (J) Peak mPFC PYR activity for each WT::PV-Cre mouse comparing Baseline (BL) (-5 to -3s before bout onset) Pre (-2 to 0s) and Post (0 to 2s) (K) Calcium traces averaged across *Mecp2* KO::PV-Cre mice during behavioral interactions with the cage-mate. Dark red line denotes average signal, shaded red denotes SEM. Time zero denotes behavioral bout onset (L) Heatmap depicting individual M *Mecp2* KO::PV-Cre mice calcium response during behavioral interactions with the cage-mate (M) Peak mPFC PYR activity for each *Mecp2* KO::PV-Cre mouse comparing Baseline (BL) (-5 to -3s before bout onset) Pre (-2 to 0s) and Post (0 to 2s) (N) Calcium traces averaged across *Mecp2* KO::PV-Cre mice during behavioral interactions with a novel conspecific. Dark red line denotes average signal, shaded red denotes SEM. Time zero denotes behavioral bout onset (O) Heatmap depicting individual calcium response during behavioral interactions with a novel conspecific of *Mecp2* KO::PV-Cre mice (P) Peak mPFC PYR activity for each *Mecp2* KO::PV-Cre mouse comparing Baseline (BL) (-5 to -3s before bout onset) Pre (-2 to 0s) and Post (0 to 2s). Data are mean ±SEM.

### 1. Validation of new 4-chamber social arena

To reduce the effort and distance needed by a tethered mouse to move between different social choices, we modified the commercially available *SocioBox* that originally has 1 central chamber surrounded by 5 side chambers to enclose different social stimuli (Krueger-Burg et al., 2016), by removing 2 of the side chambers to simplify choices for the test mouse (Suppl. Fig. 1A). All mice were first habituated to the social arena with empty side chambers (see Material and Methods), and then underwent the following behavioral testing: 1) habituation phase with empty side chambers; 2) *sociability* phase with a social object (toy mouse) and a sex-matched cage-mate in different side chambers (Suppl. Fig. 1B); and 3) *social memory* phase with a novel sex- and age-matched mouse added to the 3rd side chamber (Suppl. Fig. 1E). Behavior events were quantified as the duration of time spent within an interaction zone of 2.5cm from the perforated partition with the side chamber (Suppl. Fig. 1A), where the experimental mouse could explore a social object (toy mouse) and interact with either a sex-matched cage-mate or a sex- and age-matched novel mouse, which included whisking behaviors.

Consistent with results using a linear 3-chamber social arena (Moy et al., 2004; Nadler et al., 2004; Yang et al., 2011), WT mice spent more total time interacting with the cage-mate than exploring the toy mouse during the *sociability* phase (p=0.0227, paired t-test; n=7 mice; Suppl. Fig. 1C). Also, WT mice spent more total time interacting with the novel mouse than with the familiar cage-mate during the *social memory* phase (p=0.0143, paired t-test; n=7; Suppl. Fig. 1F). Using the total interaction times, the discrimination index of WT mice is significantly biased towards the live mouse in the *sociability* phase (Suppl. Fig. 1D), and towards the novel mouse during the *social memory* (p=0.1328, unpaired t-test; n=6; Suppl. Fig. 1G).

### 2. Fiber optic-tethered *Mecp2* KO mice show typical *sociability* but altered *social memory* in the new 4-chamber social arena

Next, we tested fiber optic-tethered WT::PV-Cre and *Mecp2* KO::PV-Cre mice with cannulas aiming at jGCaMP8m-expressing mPFC PYRs (*Camkii* promoter driving jGCaMP8f). Panels A in figures 1-4 show representative examples of mPFC sections to illustrate jGCaMP8m expression around the fiber-optic cannula (Fig. 1A, white box).

#### 2a. Sociability (toy mouse vs. cage-mate)

During the *sociability* phase, *Mecp2* KO::PV-Cre mice spent more total time interacting with the cage-mate than exploring the toy mouse (p=0.0002, paired t-test; n=11 *Mecp2* KO::PV-Cre mice; Fig. 1B), similar to WT::PV-Cre mice (p=0.0258, paired t-test; n=11 WT::PV-Cre; Fig. 1B), and the discrimination indices were comparable between genotypes (p=0.1395, unpaired t-test; n=11 each genotype; Fig. 1C). Further analyses revealed that WT::PV-Cre mice engaged in more interaction events with the cage-mate than *Mecp2* KO::PV-Cre mice (p=0.0189, 2-way ANOVA with Bonferroni’s post hoc test; n=11 per genotype; Suppl. Fig. 2A), but not with the toy mouse (p=0.0782, 2-way ANOVA with Bonferroni’s post hoc test; n=11 per genotype; Suppl. Fig. 2A). The number of toy exploration bouts was similar to that of cage-mate interactions in both WT::PV-Cre mice (p=0.2085, 2-way ANOVA with Bonferroni’s post hoc test; n=11 WT::PV-Cre) and *Mecp2* KO::PV-Cre mice (p=0.5264, 2-way ANOVA with Bonferroni’s post hoc test; n=11 *Mecp2* KO::PV-Cre; Suppl. Fig. 2A). However, each toy exploration bout and cage-mate interaction bout was longer in *Mecp2* KO::PV-Cre than in their WT::PV-Cre littermate control (toy: p=0.0253; cage-mate: p<0.0001; 2-way ANOVA, Bonferroni’s; n=11 each genotype; Suppl. Fig. 2B). Furthermore, the duration of each cage-mate interaction bout was longer than each toy interaction bout in *Mecp2* KO::PV-Cre mice (p=0.0202, 2-way ANOVA, Bonferroni’s; n=11 *Mecp2* KO), a difference absent in WT::PV-Cre mice (p=0.8169, 2-way ANOVA, Bonferroni’s; n=11 WT; Suppl. Fig. 2B).

#### 2b. Social memory (novel vs. cage-mate)

During the *social memory* phase, *Mecp2* KO::PV-Cre mice did not show the typical preference to interact more with the novel mouse than with the cage-mate, spending comparable times with either mouse (p=0.6327, paired t-test, n=10 *Mecp2* KO::PV-Cre; Fig. 2B). In contrast, WT::PV-Cre mice showed the expected preference, spending more time interacting with the novel mouse (p=0.0001, paired t-test, n=11 WT::PV-Cre; Fig. 2B), which resulted in a significant difference in their discrimination indices (p=0.0072, unpaired t-test, n=11 WT::PV-Cre vs. n=10 *Mecp2* KO::PV-Cre; Fig. 2C). Further analyses revealed that WT::PV-Cre mice engaged in more interacting bouts with the novel mouse than *Mecp2* KO::PV-Cre mice (p=0.0023, 2-way ANOVA with Bonferroni’s; n=11 per genotype; Suppl. Fig. 2C), but not with the cage-mate (p=0.1170, 2-way ANOVA with Bonferroni’s; n=11 per genotype; Suppl. Fig. 2C). Interestingly, WT::PV-Cre mice showed more interaction bouts with the novel mouse than with the cage-mate (p=0.0464, 2-way ANOVA with Bonferroni’s; n=11 WT::PV-Cre; Suppl. Fig. 2C), a difference absent in *Mecp2* KO::PV-Cre mice (p=0.6965, 2-way ANOVA with Bonferroni’s; n=11 *Mecp2* KO::PV-Cre; Suppl. Fig. 2C). The duration of individual interaction bouts with the novel mouse were longer in *Mecp2* KO::PV-Cre mice than in WT::PV-Cre (p=0.0086, 2-way ANOVA, Bonferroni’s; n=11 per genotype; Suppl. Fig. 2D), a difference also observed in the interaction bouts with the cage-mate (p=0.0001, 2-way ANOVA, Bonferroni’s; n=11 per genotype; Suppl. Fig. 2D). Furthermore, the duration of each interacting bout with the novel mouse was longer than those with the cage-mate in WT::PV-Cre mice (p=0.0287, 2-way ANOVA with Bonferroni’s; n=11 WT::PV-Cre; Suppl. Fig. 2D), a difference absent in *Mecp2* KO::PV-Cre mice (p=0.4170, 2-way ANOVA with Bonferroni’s; n=11 *Mecp2* KO::PV-Cre; Suppl. Fig. 2D).

Similar results were obtained in fiber optic-tethered WT::PV-Cre and *Mecp2* KO::PV-Cre mice with cannulas aiming at jGCaMP8m-expressing mPFC PV-INs (*Synapsin* promoter driving FLEX to express jGCaMP8f in Cre-expressing neurons); see Supplemental Results. Of note, the average velocity and total distance travelled during the entire test were smaller in *Mecp2* KO mice (velocity: p=0.0271, unpaired t-test, n=9 WT vs. n=7 *Mecp2* KO, Suppl. Fig. 2I; total distance travelled: p=0.0195 unpaired t-test, n=9 WT vs. n=7 *Mecp2* KO, Suppl. Fig. 2J), consistent with other reports of motor dysfunction in *Mecp2*-deficient mice (Samaco et al., 2013; Yue et al., 2021).

These observations demonstrate the validity of our new 4-chamber arena with fiber optic-tethered mice expressing Ca^2+^ sensors either in mPFC PYRs or PV-INs, and with implanted cannulas in the mPFC. This approach confirmed the selective deficit in *social memory* of *Mecp2* KO mice first described in untethered naive mice using a linear 3-chamber arena (Phillips et al., 2019).

### 3. The spiking activity of mPFC PYRs during social interactions is lower and more temporally spread-out in *Mecp2* KO::PV-Cre mice than in WT::PV-Cre mice

During the social behavior tasks described above, entry into the 2.5cm interaction zone was defined as time zero to synchronize simultaneously recorded fluorescent signals from the Ca^2+^ sensor jGCaMP8m expressed either in PYRs (*Camkii* promoter) or in PV-INs (Cre-dependent jGCaMP8m construct) in the mPFC of WT::PV-Cre and *Mecp2* KO::PV-Cre mice. To include those mPFC Ca^2+^ signals known to precede social interactions in similar assays (Bicks et al., 2020), we divided the time before entry into the interaction zone into a 2-second epoch to obtain *Baseline* activity (BL: -5s to -3s) and a 2-second epoch just before entry (Pre: -2s to 0s); Ca^2+^ sensor signals were analyzed for a 2 second epoch after interaction zone entry (Post: 0 to 2s). We video-recorded behavioral data at 40 fps while simultaneously recording jGCaMP8m photometry signals at 40 fps (Martianova et al., 2019; Zhang et al., 2023) (see Material and Methods). We used the maximum value (peak) of the z-scored jGCaMP8m deltaF/F trace as a proxy of both the number of neurons spiking near the fiber and the intracellular Ca^2+^ levels reached by each of those neurons. We used the area-under-the-curve (AUC) of the z-scored deltaF/F trace to estimate the temporal structure (i.e. dynamics) of neuronal population spiking near the fiber (i.e. synchronized population spiking).

#### 3a. *Sociability* (toy mouse vs. cage-mate)

In the *sociability* phase, and consistent with others (Lee et al., 2016; Liang et al., 2018), mPFC PYRs of WT::PV-Cre mice increased their spiking activity during interactions with the cage-mate, i.e. they showed larger z-scored dF/F (p=0.003 Pre vs. Post; paired t-test; n=11 WT::PV-Cre; Fig. 1H-J), while their activity remained similar prior to interaction onset (p=0.884 BL vs. Pre, paired t-test; n=11 WT::PV-Cre; Fig. 1J). In contrast, mPFC PYR activity in WT::PV-Cre mice did not change when exploring the toy mouse (p=0.088 Pre vs. Post, paired t-test; n=11 WT::PV-Cre), or entering into that interaction zone (p=0.779 BL vs. Pre, paired t-test; n=11 WT::PV-Cre; Fig. 1E-G). On the other hand, the spiking activity of mPFC PYRs in *Mecp2* KO::PV-Cre mice did not change neither during interactions with the cage-mate (p=0.355 BL vs. Pre, p=0.062 Pre vs. Post; paired t-test; n=11 *Mecp2* KO::PV-Cre; Fig. 1N-P) nor explorations of the toy mouse (p=0.263 BL vs. Pre, p=0.844 Pre vs. Post, paired t-test; n=11 *Mecp2* KO::PV-Cre; Fig. 1K-M). Similar patterns between WT::PV-Cre and *Mecp2* KO::PV-Cre were obtained when using the AUC of jGCaMP8m transients to estimate the dynamics of PYR spiking activity (Suppl. Fig. 3A-D).

These observations confirm the rapid engagement of mPFC PYRs in WT mice during interactions with a conspecific (Lee et al., 2016; Liang et al., 2018), and demonstrate a clear dysfunction of mPFC PYRs in *Mecp2* KO mice during the *sociability* phase of the social behavior task.

#### 3b. *Social memory* (novel vs. cage-mate)

In the *social memory* phase, and consistent with others (Lee et al., 2016; Liang et al., 2018), mPFC PYRs of WT::PV-Cre mice increased their spiking activity during interactions with the novel mouse (p=0.240 BL vs. Pre, p=0.005 Pre vs. Post; paired t-test; n=11 WT::PV-Cre; Fig. 2H-J). mPFC PYR activity also increased during interactions with the cage-mate (p=0.480 BL vs. Pre, p=0.017 Pre vs. Post; paired t-test; n=11 WT::PV-Cre; Fig. 2E-G). In contrast, the spiking activity of mPFC PYRs of *Mecp2* KO::PV-Cre mice did not change during interactions with either the novel mouse (p=0.081 BL vs. Pre, p=0.105 Pre vs. Post; paired t-test; n=11 *Mecp2* KO::PV-Cre; Fig. 2N-P) nor the cage-mate (p=0.874 BL vs. Pre, p=0.116 Pre vs. Post, paired t-test; n=11 *Mecp2* KO::PV-Cre; Fig. 2K-M).

Similar patterns between WT::PV-Cre and *Mecp2* KO::PV-Cre were obtained when using the AUC of jGCaMP8m transients to estimate the dynamics of PYR spiking activity (Suppl. Fig. 3E-H). Interestingly, the spiking dynamic of mPFC PYR of *Mecp2* KO::PV-Cre was heightened compared to that of WT::PV-Cre mice, but only prior to the interactions with the novel mouse (p=0.0169, unpaired t-test, n= 10 mice per genotype; Suppl. Fig. 3G,H). Of note, such spiking dynamic of mPFC PYRs in *Mecp2* KO::PV-Cre was blunted compared to that of mPFC PV-INs, but only when interacting with the cage-mate both prior to entering the interaction zone (p=0.0019) and during interactions (p=0.0067, unpaired t-test, n= 8 *Mecp2* KO::PV-Cre; Suppl. Fig. 4M,N).

These data suggest that mPFC PYRs of *Mecp2* KO mice show broader (i.e. desynchronized) spiking activity during interactions with novel mice, unlike those of WT mice, which show jGCaMP8m transients more tightly modulated during those interactions.

### 4. The spiking activity of mPFC PV-INs during social interactions is higher and lacks selectivity for the novel mouse in *Mecp2* KO::PV-Cre mice compared to that in WT::PV-Cre mice

We then injected AAVs expressing Cre-dependent jGCaMP8m (*Synapsin* promoter driving FLEX to express jGCaMP8f in Cre-expressing neurons) into the mPFC of WT::PV-Cre and *Mecp2* KO::PV-Cre mice to record spiking activity from PV-INs during the same social test.

#### 4a. *Sociability* (toy mouse vs. cage-mate)

In the *sociability* phase, and consistent with others (Bicks et al., 2020; Liu et al., 2020a; Selimbeyoglu et al., 2017; Xu et al., 2022), mPFC PV-INs of WT::PV-Cre mice increased their spiking activity during interactions with the cage-mate (p=0.0319 Pre vs. Post; paired t-test; n=8 WT::PV-Cre; Fig. 3G-I) while their activity remained similar prior to interaction onset (p=0.312 BL vs. Pre; paired t-test; n=8 WT::PV-Cre; Fig. 3I). In contrast, mPFC PV-INs activity in WT::PV-Cre mice did not change neither prior (p=0.700 BL vs. Pre; paired t-test; n=8 WT::PV-Cre) nor during exploration of the toy mouse (p=0.069 Pre vs. Post; paired t-test; n=8 WT::PV-Cre; Fig. 3D-F). On the other hand, mPFC PV-INs of *Mecp2* KO::PV-Cre mice did not show such selectivity for the live mouse: their activity increased equally during interactions with either the cage mate (p=0.501 Pre vs. Post; paired t-test; n=8 *Mecp2* KO::PV-Cre; Fig. 3M-O) or the toy mouse (p=0.437 BL vs. Pre, p=0.449 Pre vs. Post; paired t-test; n=9 *Mecp2* KO::PV-Cre; Fig. 3J-L). Of note, mPFC PV-INs of *Mecp2* KO::PV-Cre mice increased their activity just before entry into the interaction zone facing the cage-mate (p=0.015 BL vs. Pre; Fig. 3O).

**Figure 3:**
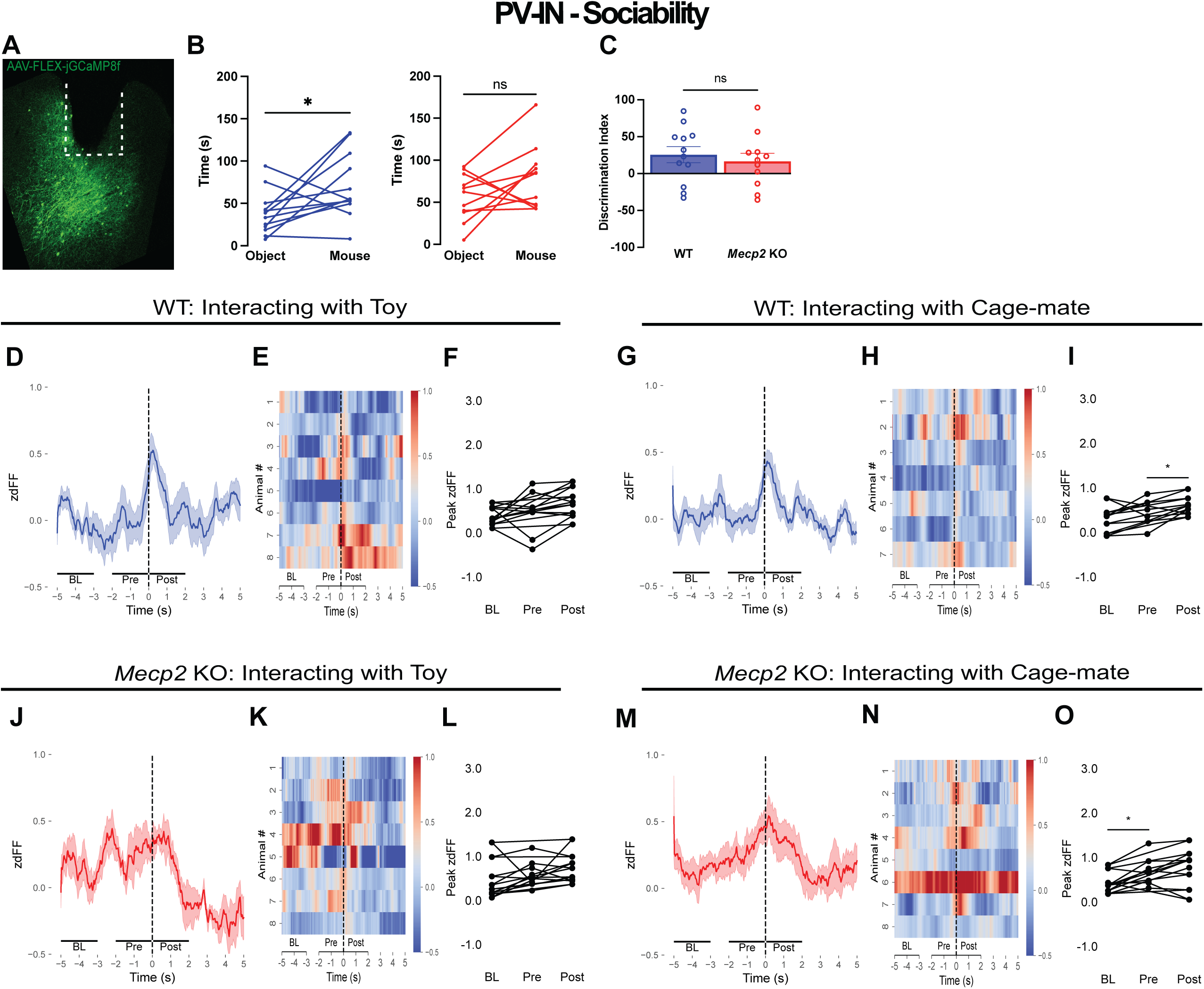
The spiking activity of mPFC PV-INs during social interactions is higher in *Mecp2* KO::PV-Cre mice compared to that in WT::PV-Cre mice. (A) Representative histological image of GCaMP8f and fiber placement (dotted line) in the mPFC. Scale bar, 100 µm. (B) Time spent interacting with the object (toy mouse) or cage-mate during the sociability trial of the 4-Chamber social interaction test for WT::PV-Cre(left) and *Mecp2* KO::PV-Cre mice (right) (C) Discrimination indices of sociability trial of WT::PV-Cre and *Mecp2* KO::PV-Cre mice (D) Calcium traces averaged across WT::PV-Cre animals during behavioral interactions with the toy mouse. Dark blue line denotes average signal, shaded blue denotes SEM. Time zero denotes behavioral bout onset (E) Heatmap depicting individual calcium response during behavioral interactions with the toy mouse of WT::PV-Cre mice (F) Peak mPFC PV-IN activity for each WT::PV-Cre mouse comparing Baseline (BL) (-5 to -3s before bout onset) Pre (-2 to 0s) and Post (0 to 2s) (G) Calcium traces averaged across WT::PV-Cre animals during behavioral interactions with the cage-mate. Dark blue line denotes average signal, shaded blue denotes SEM. Time zero denotes behavioral bout onset (H) Heatmap depicting individual calcium response during behavioral interactions with the cage-mate of WT::PV-Cre mice (I) Peak mPFC PV-IN activity for each WT::PV-Cre mouse comparing Baseline (BL) (-5 to -3s before bout onset) Pre (-2 to 0s) and Post (0 to 2s) (J) Calcium traces averaged across *Mecp2* KO::PV-Cre mice during behavioral interactions with the toy mouse. Dark redline denotes average signal, shaded red denotes SEM. Time zero denotes behavioral bout onset (K) Heatmap depicting individual *Mecp2* KO::PV-Cre mice calcium response during behavioral interactions with the toy mouse (L) Peak mPFC PV-IN activity for each *Mecp2* KO::PV-Cre mouse comparing Baseline (BL) (-5 to -3s before bout onset) Pre (-2 to 0s) and Post (0 to 2s) (M) Calcium traces averaged across *Mecp2* KO::PV-Cre mice during behavioral interactions with he cage-mate Dark red line denotes average signal, shaded red denotes SEM. Time zero denotes behavioral bout onset (N) Heatmap depicting individual calcium response during behavioral interactions with he cage-mate of *Mecp2* KO::PV-Cre mice (O) Peak mPFC PV-IN activity for each *Mecp2* KO::PV-Cre mouse comparing Baseline (BL) (-5 to -3s before bout onset) Pre (-2 to 0s) and Post (0 to 2s). Data are mean ±SEM.

**Figure 4:**
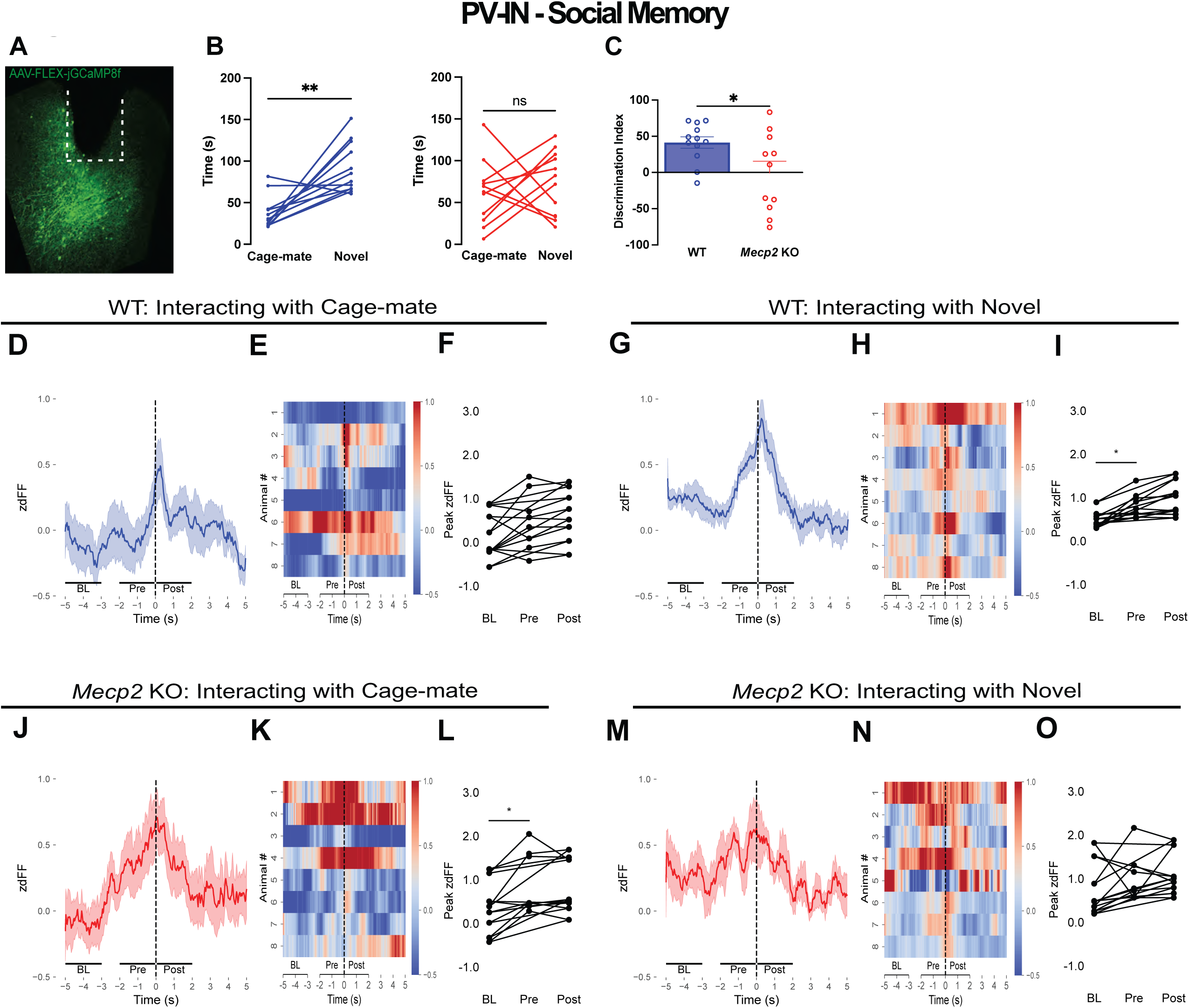
The spiking activity of mPFC PV-INs during social interactions lacks selectivity for the novel mouse in *Mecp2* KO::PV-Cre mice compared to that in WT::PV-Cre mice. (A) Representative histological image of GCaMP8f and fiber placement (dotted line) in the mPFC. Scale bar, 100 µm. (B) Time spent interacting with the cage-mate or novel mouse during the social memory trial of the 4-Chamber social interaction test for WT::PV-Cre (left) and *Mecp2* KO::PV-Cre mice (right) (C) Discrimination indices of the social memory trial of *Mecp2* KO::PV-Cre and WT::PV-Cre mice (D) Calcium traces averaged across WT::PV-Cre animals during behavioral interactions with the cage-mate. Dark blue line denotes average signal, shaded blue denotes SEM. Time zero denotes behavioral bout onset (E) Heatmap depicting individual calcium response in WT::PV-Cre mice during behavioral interactions with the cage-mate (F) Peak mPFC PV-IN activity for each WT::PV-Cre mouse comparing Baseline (BL) (-5 to -3s before bout onset) Pre (-2 to 0s) and Post (0 to 2s) (G) Calcium traces averaged across WT::PV-Cre animals during behavioral interactions with the novel mouse. Dark blue line denotes average signal, shaded blue denotes SEM. Time zero denotes behavioral bout onset (H) Heatmap depicting individual calcium response in WT::PV-Cre mice during behavioral interactions with the novel mouse (I) Peak mPFC PV-IN activity for each WT::PV-Cre mouse comparing Baseline (BL) (-5 to -3s before bout onset) Pre (-2 to 0s) and Post (0 to 2s) (J) Calcium traces averaged across *Mecp2* KO::PV-Cre mice during behavioral interactions with the cage-mate. Dark red line denotes average signal, shaded red denotes SEM. Time zero denotes behavioral bout onset (K) Heatmap depicting individual calcium response in *Mecp2* KO::PV-Cre mice during behavioral interactions with the cage-mate (L) Peak mPFC PV-IN activity for each *Mecp2* KO::PV-Cre mouse comparing Baseline (BL) (-5 to -3s before bout onset) Pre (-2 to 0s) and Post (0 to 2s) (M) Calcium traces averaged across *Mecp2* KO::PV-Cre mice during behavioral interactions with a novel conspecific. Dark red line denotes average signal, shaded red denotes SEM. Time zero denotes behavioral bout onset (N) Heatmap depicting individual calcium response of *Mecp2* KO::PV-Cre mice during behavioral interactions with the novel mouse (O) Peak mPFC PV-IN activity for each *Mecp2* KO::PV-Cre mouse comparing Baseline (BL) (-5 to -3s before bout onset) Pre (-2 to 0s) and Post (0 to 2s). Data are mean ±SEM.

Similar patterns between WT::PV-Cre and *Mecp2* KO::PV-Cre were obtained when using the AUC of jGCaMP8m transients to estimate the dynamics of PV-IN spiking activity (Suppl. Fig. 3I-L).

These data suggest that mPFC PV-INs of *Mecp2* KO::PV-Cre mice lack the typical selective increase of activity during the *sociability* phase (toy vs. cage-mate) shown by those of WT::PV-Cre mice, with an atypical increase of activity when exploring the toy mouse. In addition, their activity precedes and spreads more than that of WT::PV-Cre mice mice during interactions with the cage mate.

#### 4b. *Social memory* (novel vs. cage-mate)

In the *social memory* phase, the spiking activity of mPFC PV-INs in WT::PV-Cre mice increased during interactions only with the novel mouse (p=0.0079 BL vs. Pre, p=0.0727 Pre vs. Post; paired t-test; n=8 WT::PV-Cre; Fig. 4G-I), but not with the cage-mate (p=0.086 Pre vs. Post, paired t-test; n=8 WT::PV-Cre; Fig. 4D-F). Consistent with others (Bicks et al., 2020), the activity of mPFC PV-INs increased from the 2-sec BL to the epoch prior to interaction zone entry (p=0.0079 BL vs. Pre) without further increases upon entry into the interaction zone, but only for interactions with the novel mouse (p=0.0727 Pre vs. Post, paired t-test; n=8 WT::PV-Cre; Fig. 4I). In contrast, the spiking activity of mPFC PV-INs of *Mecp2* KO::PV-Cre mice did not precede nor change during interactions with the novel mouse (p=0.431 BL vs. Pre, p=0.576 Pre vs. Post; paired t-test; n=8 *Mecp2* KO::PV-Cre; Fig. 4M-O). Also distinct from WT::PV-Cre mice, their activity increased from the 2-sec BL to the epoch prior to interaction zone entry (p=0.0074 BL vs. Pre) without further increases upon entry into the interaction zone, but only for interactions with the cage-mate (p=0.368 Pre vs. Post; paired t-test; n=8 *Mecp2* KO::PV-Cre; Fig. 4J-L).

Similar patterns between WT::PV-Cre and *Mecp2* KO::PV-Cre were obtained when using the AUC of jGCaMP8m transients to estimate the dynamics of PV-IN spiking activity (Suppl. Fig. 3M-P). Interestingly, the spiking dynamics of mPFC PV-INs was heightened compared to that of PYRs in *Mecp2* KO::PV-Cre mice, both prior (p=0.0019) and during interactions with the cage-mate (p=0.0067, unpaired t-test, n=8 *Mecp2* KO::PV-Cre; Suppl. Fig. 4M,N).

These data suggest that mPFC PV-INs of *Mecp2* KO::PV-Cre mice lack the typical selective increase of activity during the *social memory* phase (cage-mate vs. novel) shown by those of WT::PV-Cre mice, with an atypical increase of activity when exploring the cage-mate. In addition, their activity precedes and spreads more than that of WT::PV-Cre mice during interactions with the cage mate.

In summary, our findings reveal distinct neural responses in WT::PV-Cre mice compared to *Mecp2* KO::PV-Cre mice during social interactions. While WT::PV-Cre mice exhibit behaviorally-relevant responses to novel conspecifics, indicating intact social memory processing, *Mecp2* KO::PV-Cre mice display altered responses, suggesting deficits in social memory. Specifically, *Mecp2* KO::PV-Cre mice show unresponsive PV-IN activity during interaction bouts with novel mice, whereas they exhibit responsive PV-IN signals during interactions with familiar conspecifics, which are broader and wider compared to WT::PV-Cre mice.

### 5. Dual-color fiber photometry recordings show similar patterns to those from single-color recordings in both mPFC PYRs and PV-INs

Since both glutamatergic PYRs and GABAergic INs in the mPFC increase their spiking activity during social interactions (Lee et al., 2016; Liang et al., 2018; Liu et al., 2020a; Selimbeyoglu et al., 2017; Xu et al., 2022; Zhao et al., 2022) (see Fig. 1-4), we also performed simultaneous dual-color Ca^2+^ fiber photometry from mPFC PYRs and PV-INs in the same mouse using *Camkii*-promoter-driven green-shifted jGCaMP8m for PYRs and Cre-dependent red-shifted jRGECO for PV-INs (Akerboom et al., 2013; Suthard et al., 2024) in *Mecp2* KO::PV-Cre mice and their WT::PV-Cre littermates (see Material and Methods).

First, we confirmed that *Mecp2* KO::PV-Cre mice injected with a mixture of AAV1-*Camkiia*-jGCaMP8m and AAV1-*Syn*-FLEX-jRGECO1a, and with fiber optic cannulas in the mPFC, show the typical *sociability* but altered *social memory* compared to their WT::PV-Cre littermate controls (see Supplemental Results), as in our previous results here (see Fig. 1-4), and in untethered naive mice using a linear 3-chamber arena (Phillips et al., 2019).

#### 5a. *Sociability* (toy mouse vs. cage-mate)

In the *sociability* phase, and consistent with our single-color photometry results (see Fig. 1), mPFC PYRs from WT::PV-Cre mice increased their spiking activity prior to the onset of interactions with the cage-mate (p=0.005 BL vs. Pre, paired t-test; n=7 WT::PV-Cre; Fig. 5F-H), and their activity remained similar following entry into the interaction zone (p=0.052 Pre vs. Post, paired t-test; n=7 WT mice; Fig. 5F-H). Similarly, both mPFC PYRs and PV-INs increased their activity prior to toy exploration (PYR: p=0.016 BL vs. Pre, paired t-test; n=7 WT::PV-Cre; Fig. 5A-C; PV-IN: p=0.005 BL vs. Pre, paired t-test; n=7 WT::PV-Cre; Fig. 5A,D,E), and their activity remained unchanged following entry into the interaction zone (PYR: p=0.119 Pre vs. Post, paired t-test; n=7 WT::PV-Cre; Fig. 5A-C; PV-IN: p=0.082 Pre vs. Post, paired t-test; n=7 WT::PV-Cre; Fig. 5A,D,E). On the other hand, the spiking activity of mPFC PYRs in *Mecp2* KO::PV-Cre mice did not change neither during interactions with the cage-mate (p=0.356 BL vs. Pre, p=0.151 Pre vs. Post, paired t-test; n=5 *Mecp2* KO::PV-Cre; Fig. 5P-R) nor during toy exploration (p=0.913 BL vs. Pre, p=0.273 Pre vs. Post, paired t-test; n=5 *Mecp2* KO::PV-Cre; Fig. 5K-M). Interestingly, mPFC PV-IN activity in *Mecp2* KO::PV-Cre mice increased only prior to interactions with the cage-mate (p=0.0165 BL vs. Pre, paired t-test; n=5 *Mecp2* KO::PV-Cre; Fig. 5P,S,T), while their activity remained unchanged following entry into the interaction zone (p=0.276 BL vs. Pre, paired t-test; n=5 *Mecp2* KO::PV-Cre; Fig. 5P,S,T). PV-IN activity in *Mecp2* KO::PV-Cre mice also remained unchanged during toy explorations (p=0.621 BL vs. Pre, p=0.241 Pre vs. Post, paired t-test; n=5 *Mecp2* KO::PV-Cre; Fig. 5K,N,O).

**Figure 5:**
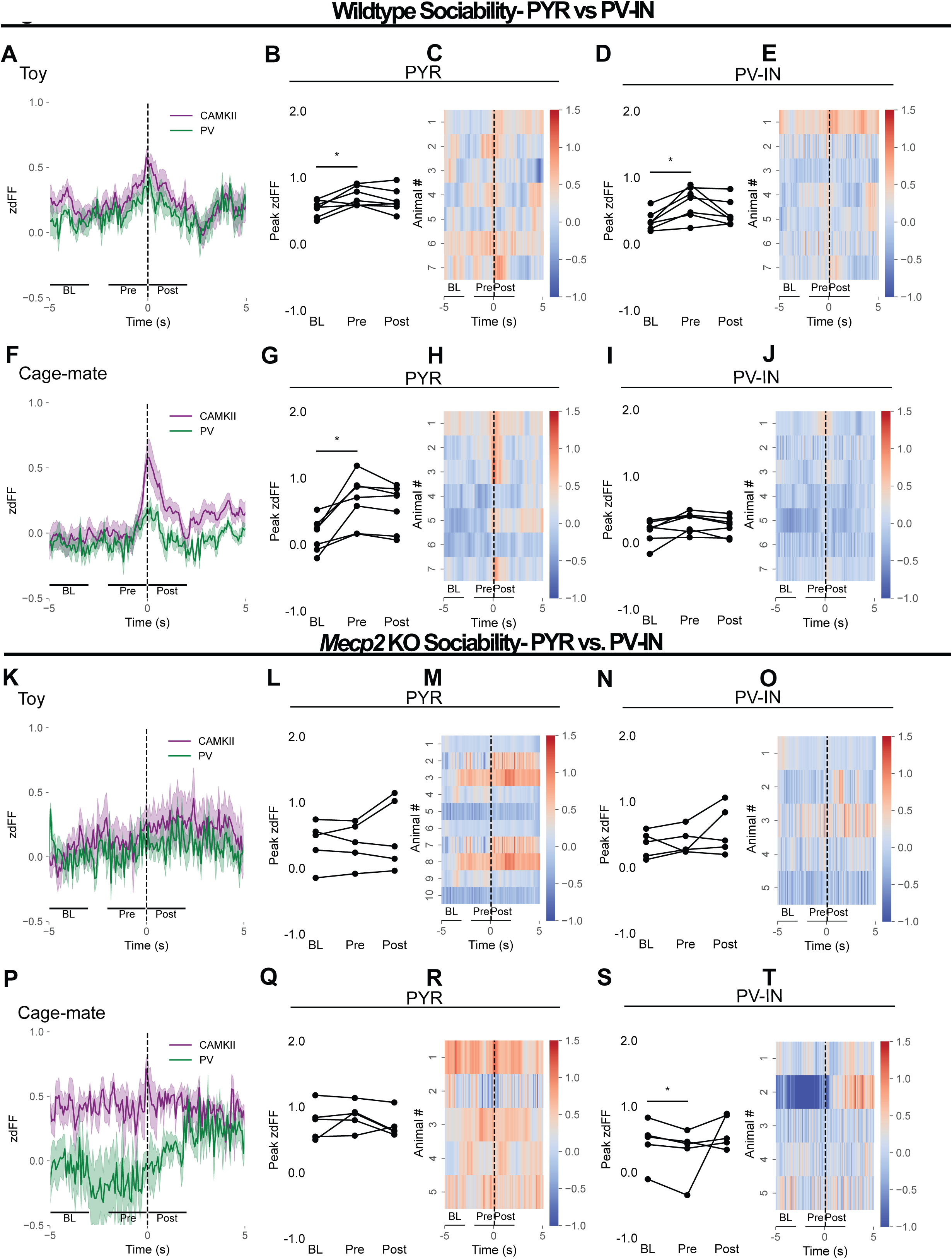
Dual color fiber photometry recordings demonstrate similar trends to single color GCaMP recordings in both PYRs and PV-INs: Sociability. (A) Simultaneous recording of *Camkii*-GCaMP8m (PYR) and *FLEX*-jRGECO (PV-IN) in WT::PV-Cre mice calcium traces averaged across animals during behavioral interactions with the toy mouse. Dark purple line denotes average signal from PYRs (CamKII), shaded purple denotes SEM. Dark green line denotes average signal from PV-INs, shaded green denotes SEM. Time zero denotes behavioral bout onset (B) Peak mPFC PYR activity for each WT::PV-Cre mouse comparing Baseline (BL) (-5 to -3s before bout onset) Pre (-2 to 0s) and Post (0 to 2s) (C) Heatmap depicting PYR calcium response in WT::PV-Cre mice during behavioral interactions with the toy mouse (D) Peak mPFC PV-IN activity for each WT::PV-Cre mouse comparing Baseline (BL) (-5 to -3s before bout onset) Pre (-2 to 0s) and Post (0 to 2s) (E) Heatmap depicting PV calcium response in WT::PV-Cre mice during behavioral interactions with the toy mouse (F) Simultaneous recording of CaMKII-GCaMP8m (PYR) and FLEX-jRGECO (PV) in WT::PV-Cre mice calcium traces averaged across mice during behavioral interactions with the cage-mate. Dark purple line denotes average signal from PYR, shaded purple denotes SEM. Dark green line denotes average signal from PV-INs, shaded green denotes SEM. Time zero denotes behavioral bout onset (G) Peak mPFC PYR activity for each WT::PV-Cre mouse comparing Baseline (BL) (-5 to -3s before bout onset) Pre (-2 to 0s) and Post (0 to 2s) (H) Heatmap depicting PYR calcium response in WT::PV-Cre mice during behavioral interactions with the cage-mate (I) Peak mPFC PV-IN activity for each WT::PV-Cre mouse comparing Baseline (BL) (-5 to -3s before bout onset) Pre (-2 to 0s) and Post (0 to 2s) (J) Heatmap depicting PV calcium response in WT::PV-Cre mice during behavioral interactions with the cage-mate (K) Simultaneous recording of CaMKII-GCaMP8m (PYR) and FLEX-jRGECO (PV) in calcium traces averaged across *Mecp2* KO::PV-Cre mice during behavioral interactions with the toy mouse. Dark purple line denotes average signal from PYR, shaded purple denotes SEM. Dark green line denotes average signal from PV-INs, shaded green denotes SEM. Time zero denotes behavioral bout onset (L) Peak mPFC PYR activity for each *Mecp2* KO::PV-Cre mouse comparing Baseline (BL) (-5 to -3s before bout onset) Pre (-2 to 0s) and Post (0 to 2s) (M) Heatmap depicting PYR calcium response in *Mecp2* KO::PV-Cre mice during behavioral interactions with the toy mouse (N) Peak mPFC PV-IN activity for each *Mecp2* KO::PV-Cre mouse comparing Baseline (BL) (-5 to -3s before bout onset) Pre (-2 to 0s) and Post (0 to 2s) (O) Heatmap depicting PV calcium response in *Mecp2* KO::PV-Cre mice during behavioral interactions with the toy mouse (P) Simultaneous recording of CaMKII-GCaMP8m (PYR) and FLEX-jRGECO (PV) in *Mecp2* KO::PV-Cre mice calcium traces averaged across mice during behavioral interactions with the cage-mate. Dark purple line denotes average signal from PYR, shaded purple denotes SEM. Dark green line denotes average signal from PV-INs, shaded green denotes SEM. Time zero denotes behavioral bout onset (Q) Peak mPFC PYR activity for each *Mecp2* KO::PV-Cre mouse comparing Baseline (BL) (-5 to -3s before bout onset) Pre (-2 to 0s) and Post (0 to 2s) (R) Heatmap depicting PYR calcium response in *Mecp2* KO::PV-Cre mice during behavioral interactions with the cage-mate (S) Peak mPFC PV-IN activity for each *Mecp2* KO::PV-Cre mouse comparing Baseline (BL) (-5 to -3s before bout onset) Pre (-2 to 0s) and Post (0 to 2s) (T) Heatmap depicting PV calcium response in *Mecp2* KO::PV-Cre mice during behavioral interactions with the cage-mate

**Figure 6:**
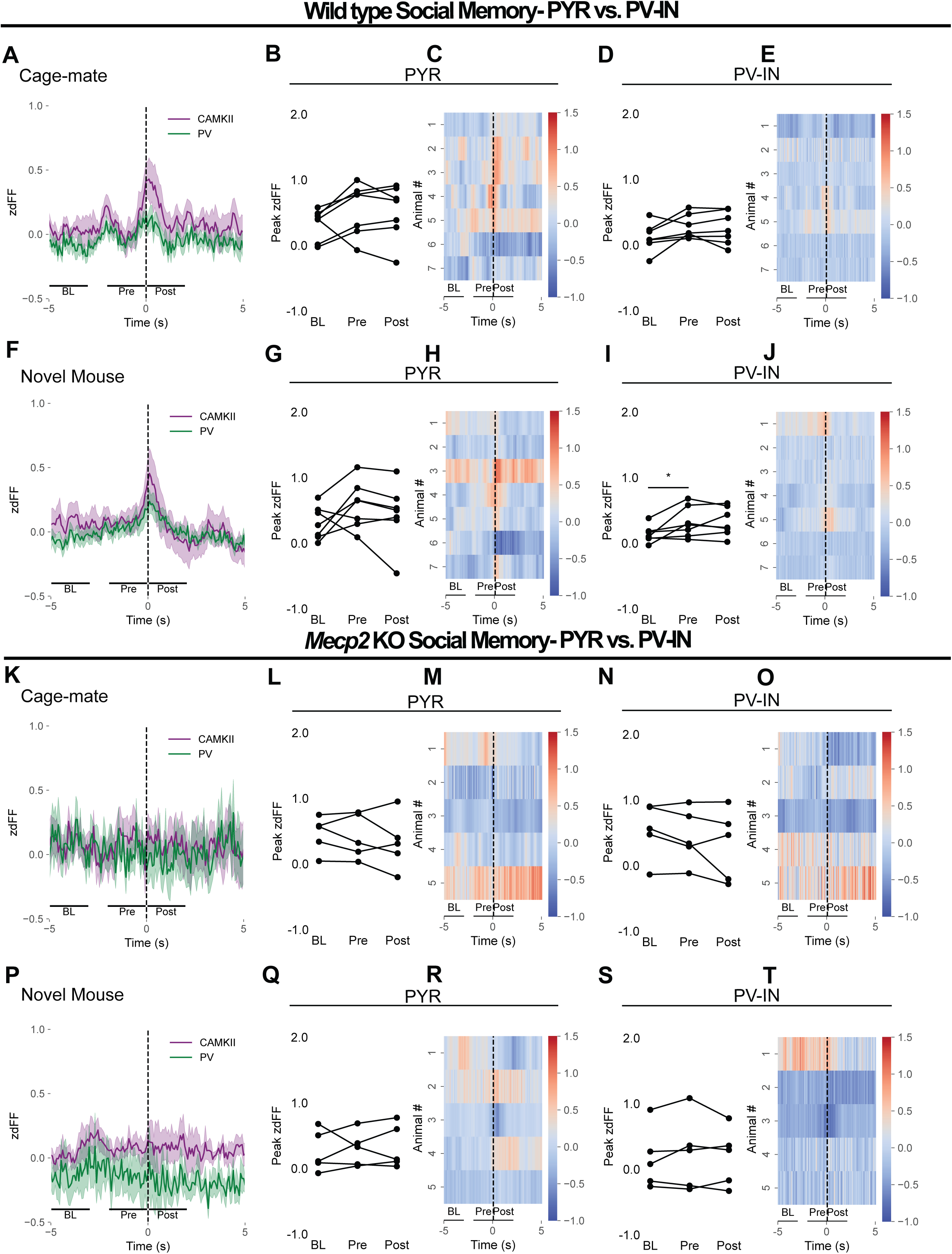
Dual color fiber photometry recordings demonstrate similar trends to single color GCaMP recordings in both PYRs and PV-INs: Social Memory. (A) Simultaneous recording of *Camkii*-GCaMP8m (PYR) and *FLEX*-jRGECO (PV-IN) in WT::PV-Cre mice calcium traces averaged across animals during behavioral interactions with the cage-mate. Dark purple line denotes average signal from PYRs (CamKII), shaded purple denotes SEM. Dark green line denotes average signal from PV-INs, shaded green denotes SEM. Time zero denotes behavioral bout onset (B) Peak mPFC PYR activity for each WT::PV-Cre mouse comparing Baseline (BL) (-5 to -3s before bout onset) Pre (-2 to 0s) and Post (0 to 2s) (C) Heatmap depicting PYR calcium response in WT::PV-Cre mice during behavioral interactions with the cage-mate (D) Peak mPFC PV-IN activity for each WT::PV-Cre mouse comparing Baseline (BL) (-5 to -3s before bout onset) Pre (-2 to 0s) and Post (0 to 2s) (E) Heatmap depicting PV calcium response in WT::PV-Cre mice during behavioral interactions with the cage-mate (F) Simultaneous recording of CaMKII-GCaMP8m (PYR) and FLEX-jRGECO (PV) in WT::PV-Cre mice calcium traces averaged across mice during behavioral interactions with the novel mouse. Dark purple line denotes average signal from PYR, shaded purple denotes SEM. Dark green line denotes average signal from PV-INs, shaded green denotes SEM. Time zero denotes behavioral bout onset (G) Peak mPFC PYR activity for each WT::PV-Cre mouse comparing Baseline (BL) (-5 to -3s before bout onset) Pre (-2 to 0s) and Post (0 to 2s) (H) Heatmap depicting PYR calcium response in WT::PV-Cre mice during behavioral interactions with the novel mouse (I) Peak mPFC PV-IN activity for each WT::PV-Cre mouse comparing Baseline (BL) (-5 to -3s before bout onset) Pre (-2 to 0s) and Post (0 to 2s) (J) Heatmap depicting PV calcium response in WT::PV-Cre mice during behavioral interactions with the novel mouse (K) Simultaneous recording of CaMKII-GCaMP8m (PYR) and FLEX-jRGECO (PV) in calcium traces averaged across *Mecp2* KO::PV-Cre mice during behavioral interactions with the cage-mate. Dark purple line denotes average signal from PYR, shaded purple denotes SEM. Dark green line denotes average signal from PV-INs, shaded green denotes SEM. Time zero denotes behavioral bout onset (L) Peak mPFC PYR activity for each *Mecp2* KO::PV-Cre mouse comparing Baseline (BL) (-5 to -3s before bout onset) Pre (-2 to 0s) and Post (0 to 2s) (M) Heatmap depicting PYR calcium response in *Mecp2* KO::PV-Cre mice during behavioral interactions with the cage-mate (N) Peak mPFC PV-IN activity for each *Mecp2* KO::PV-Cre mouse comparing Baseline (BL) (-5 to -3s before bout onset) Pre (-2 to 0s) and Post (0 to 2s) (O) Heatmap depicting PV calcium response in *Mecp2* KO::PV-Cre mice during behavioral interactions with the cage-mate (P) Simultaneous recording of CaMKII-GCaMP8m (PYR) and FLEX-jRGECO (PV) in *Mecp2* KO::PV-Cre mice calcium traces averaged across mice during behavioral interactions with the novel mouse. Dark purple line denotes average signal from PYR, shaded purple denotes SEM. Dark green line denotes average signal from PV-INs, shaded green denotes SEM. Time zero denotes behavioral bout onset (Q) Peak mPFC PYR activity for each *Mecp2* KO::PV-Cre mouse comparing Baseline (BL) (-5 to -3s before bout onset) Pre (-2 to 0s) and Post (0 to 2s) (R) Heatmap depicting PYR calcium response in *Mecp2* KO::PV-Cre mice during behavioral interactions with the novel mouse (S) Peak mPFC PV-IN activity for each *Mecp2* KO::PV-Cre mouse comparing Baseline (BL) (-5 to -3s before bout onset) Pre (-2 to 0s) and Post (0 to 2s) (T) Heatmap depicting PV calcium response in *Mecp2* KO::PV-Cre mice during behavioral interactions with the novel mouse

#### 5b. *Social memory* (novel vs. cage-mate)

In the *social memory* phase, the spiking activity of mPFC PYRs of WT::PV-Cre mice did not change during interactions with either the novel mouse (p=0.126 BL vs. Pre, p=0.120 Pre vs. Post, paired t-test; n=7 WT::PV-Cre; Fig. 6F-H) nor with the cage-mate (p=0.127 BL vs. Pre, p=0.581 Pre vs. Post, paired t-test; n=7 WT::PV-Cre; Fig. 6A-C). In contrast, mPFC PV-IN activity increased in WT::PV-Cre prior to the onset of interactions with the novel mouse (p=0.047 BL vs. Pre, paired t-test; n=7 WT::PV-Cre; Fig. 6F,I,J), while PV-IN activity remained unchanged following entry into the interaction zone (p=0.881 Pre vs. Post, paired t-test; n=7 WT::PV-Cre; Fig. 6F,I,J). However, mPFC PV-IN activity did not change during interactions with the cage-mate (p=0.076 BL vs. Pre, p=0.707 Pre vs. Post, paired t-test; n=7 WT::PV-Cre; Fig. 7A,D,E). In contrast, PYR activity in *Mecp2* KO::PV-Cre mice did not change during interactions neither with the novel mouse (p=0.739 BL vs. Pre, p=0.680 Pre vs. Post, paired t-test; n=7 WT::PV-Cre; Fig. 6P-R), nor with the cage-mate (p=0.640 BL vs. Pre, p=0.411 Pre vs. Post, paired t-test; n=7 WT::PV-Cre; Fig. 6K-M). Similarly, PV-IN activity in *Mecp2* KO::PV-Cre mice remained unchanged following entry into the interaction zones for the novel mouse (p=0.333 BL vs. Pre, p=0.541 Pre vs. Post, paired t-test; n=7 WT::PV-Cre; Fig. 6P,S,T) and the cage-mate (p=0.176 BL vs. Pre, p=0.339 Pre vs. Post, paired t-test; n=7 WT::PV-Cre; Fig. 6K,N,O).

Taken altogether, these results indicate that, while both mPFC PYRs and PV-INs of WT::PV-Cre mice increase their spiking activity during social interaction with novel mice, those in *Mecp2* KO::PV-Cre mice lack such behavioral modulation of spiking activity. These observations suggest atypically heightened inhibition and impaired excitation in the mPFC network of *Mecp2* KO::PV-Cre mice during social interactions, potentially driving their deficit in social memory.

## DISCUSSION

Here, we report our findings of spiking activity from glutamatergic excitatory pyramidal neurons (PYRs) and parvalbumin (PV)-expressing GABAergic inhibitory interneurons (PV-INs) in the mPFC of mice performing social behaviors in a new 4-chamber arena. We performed single-color photometry of the green-shifted Ca^2+^ sensor jGCaMP8 expressed in either PYRs or PV-INs of different mice, as well as dual-color photometry of jGCaMP8m expressed in PYRs and red-shifted jRGECO expressed in PV-INs. We compared the spiking activity of mPFC PYRs and PV-INs in WT::PV-Cre mice during exploration of a toy mouse, a novel mouse, and a familiar cage-mate, to the activity of those neuronal cell types in *Mecp2* KO::PV-Cre mice, an experimental model for the monogenic autism disorder Rett syndrome.

We first confirmed that our new 4-chamber social assay (modified from Krueger-Burg et al., 2016) yields results in untethered WT mice that are similar to those obtained with the classical linear 3-chamber social arena (Moy et al., 2004). Using the new 4-chamber social assay, we also confirmed the selective deficit in *social memory* of *Mecp2* KO mice first described in untethered naive mice using a linear 3-chamber arena (Phillips et al., 2019). We recorded spiking activity from mPFC neurons during a social task because of its role in social motivation and recognition in the human and rodent brain (Kim et al., 2015). We focused on PYRs and PV-INs in the mPFC due to their established roles in social behaviors and maintaining a proper excitation-inhibition balance at the network level (Bicks et al., 2020; Liang et al., 2018; Sun et al., 2020; Yizhar et al., 2011). We found that most mPFC PYRs in WT::PV-Cre mice increase their spiking activity during explorations of a familiar cage-mate mouse and a novel mouse. In contrast, most mPFC PYRs in *Mecp2* KO::PV-Cre mice did not change their activity during toy mouse exploration or interactions with a familiar mouse. Furthermore, the temporal spread of their activity during interactions with a novel mouse suggests longer desynchronized spiking epochs compared to the briefer and synchronous patterns of mPFC PYRs in WT::PV-Cre mice. Such atypical spiking pattern may reflect and/or originate from the network hypoactivity of the mPFC in *Mecp2* KO::PV-Cre mice, reported to result from either fewer excitatory inputs between PYRs (as in primary somatosensory cortex and mPFC) (Dani et al., 2005; Sceniak et al., 2016) or more abundant inhibitory inputs onto PYRs, as in primary visual cortex (Durand et al., 2012). Supporting this notion, *in vivo* Ca^2+^ imaging from mPFC PYRs in female *Mecp2* heterozygous mice carrying head-mounted miniscopes also revealed a lower spiking frequency during social interactions than in WT mice (Xu et al., 2022).

mPFC PV-INs in WT::PV-Cre mice increase their spiking activity during the first exposure to a familiar cage-mate mouse during the *sociability* phase, but not the second exposure during the *social memory* phase and during explorations of a novel mouse. In contrast, mPFC PV-INs in *Mecp2* KO::PV-Cre mice increase their activity prior to onset of interactions with a familiar cage-mate mouse. Moreover, the temporal spread of their activity during interactions with a cage-mate mouse suggests longer desynchronized spiking patterns compared to the activity of PV-INs in WT::PV-Cre mice, as well as PYR in *Mecp2* KO::PV-Cre mice. In addition, the activity of PV-INs of *Mecp2* KO::PV-Cre mice remains unaltered during interactions with a novel mouse, which may result from altered inputs from ventral hippocampal projections known to contribute to social memory (Phillips et al., 2019; Sun et al., 2020).

Excitatory projections from the ventral hippocampus to the mPFC, known to modulate social memory (Phillips et al., 2019), may propagate the hyperactivity of the hippocampal network in *Mecp2* KO mice (Calfa et al., 2011; Calfa et al., 2015; D’Cruz et al., 2010; Li et al., 2016; Wither et al., 2018; Zhang et al., 2008) to the mPFC microcircuit, further disrupting its network imbalance (Sceniak et al., 2016). Indeed, chemogenetic inhibition of mPFC-projecting ventral CA1 PYRs restored social memory preference in *Mecp2* KO mice (Phillips et al., 2019). Consistently, stimulation of mPFC PV-INs or inhibition of VIP-INs that synapse onto them also restored social memory (Sun et al., 2020). On the other hand, inhibition of mPFC PV-INs during a social assay increased PYR activity and improved social discrimination in female *Mecp2* heterozygous mice (Xu et al., 2022), suggesting that modulation of mPFC inhibition can partially restore proper mPFC function also in female Rett mice, whose brains are a mosaic of MeCP2-expressing and MeCP2-lacking neurons due to random X-chromosome inactivation (Braunschweig et al., 2004). Together, these studies support the notion that proper network balance in the mPFC is essential for social memory, and its alteration in Rett mice contributes to their social behavior deficits.

In addition to characterizing spiking activity of mPFC PYRs and PV-INs in different mice performing the same social behavior test, we also recorded simultaneously from them in the same mouse by dual-color fiber photometry of Ca^2+^ sensors with non-overlapping excitation and emission spectra (Akerboom et al., 2013; Suthard et al., 2024). These experiments confirmed that mPFC PYRs and PV-INs of WT::PV-Cre mice increase their spiking activity <u>prior</u> to both toy exploration and cage-mate interactions in the *sociability* phase, while PV-INs increase their activity prior to interactions with a novel mouse in the *social memory* phase. In contrast, mPFC PV-INs of *Mecp2* KO::PV-Cre mice increase their activity only <u>during</u> interactions with a familiar cage-mate, without any changes during toy exploration or interactions with a novel mouse. Similarly, mPFC PYRs of *Mecp2* KO::PV-Cre mice lacked changes in activity during toy exploration or interactions with a cage-mate or a novel mouse, further confirming observations from single-color fiber photometry recordings of mPFC PYRs and PV-INs in different mice.

A limitation of the approach we followed here is that photometry of ‘bulk’ fluorescence signals from Ca^2+^ sensors reports population responses, akin multiunit extracellular recordings, and not all mPFC neurons change their spiking activity during social interactions, with different PYRs that either increase or decrease their activity (Liang et al., 2018; Zhao et al., 2022). The large difference in the numbers and spatial distribution of PYRs and PV-INs in the mPFC further complicates the interpretation of population Ca^2+^ photometry recordings. Another limitation is that the red-shifted jRGECO Ca^2+^ sensor is reported to ‘photoswitch’ when illuminated with blue light, which was necessary to excite jGCaMP8 during dual-color photometry recordings (Akerboom et al., 2013; Dana et al., 2016; Shaner et al., 2008; Shen et al., 2018).

Collectively, we found that the dynamic interplay of PYRs and PV-INs in the mPFC of WT mice is altered in *Mecp2* KO, which underscores the critical role of proper mPFC network balance for proper social behaviors.

## AUTHOR CONTRIBUTIONS

DM performed experiments, analyzed data, and wrote the manuscript; LP wrote code and performed data analyses; WL designed custom equipment, performed experiments, and analyzed data; LP-M designed experiments, analyzed data, and edited the manuscript.

## ACKNOWLEDGEMENTS

We thank Mr. Cesar Acevedo-Triana and Mr. Chang Li for helpful discussions, and Dr. Kirstie Cummings (UAB) and Dr. Sofia Beas (UAB) for discussions. This work was supported by NIH grants MH-118563-01 (LP-M), MH-118563-04-S1 (DM), and T32-NS061788 (DM).

