## Supplemental Figure Legends for "Altered activity of mPFC pyramidal neurons and parvalbumin-expressing interneurons during social interactions in a *Mecp2* mouse model for Rett syndrome"

**Medeiros et al., 2024 Supplemental Figure Legends**

**Supplemental Figure 1: A new 4-chamber social interaction assay for assessing social behaviors**

(A) Representative image (upper) and schematic (lower) of the new 4-Chamber social behavior apparatus
(B) Schematic of 4-Chamber social interaction apparatus during the sociability trial with simultaneous fiber photometry recording (left) and representative heatmap of sociability trial (right)
(C) Time spent interacting with the toy object, or cage-mate during the sociability trial
(D) Discrimination index of sociability trial in wild type male mice
(E) Schematic of 4-Chamber apparatus during the social memory trial with simultaneous fiber photometry recording (left) and representative heatmap of social memory trial (right)
(F) Time spent interacting with the familiar conspecific or novel conspecific during the social memory trial
(G) Discrimination index of social memory trial in wild type male mice. Data are mean ±SEM

**Supplemental Figure 2: Reduced motor movement of *Mecp2* KO::PV-Cre mice in the 4-Chamber social assay**

(A) Number of interaction bouts with the toy object or cage-mate during the sociability trial of the 4-Chamber social interaction test for WT::PV-Cre and *Mecp2* KO::PV-Cre mice injected with *Camkii*-GCAMP8m.
(B) Duration of interaction bouts with the object or cage-mate during the sociability trial of the 4-Chamber social interaction test for WT::PV-Cre and *Mecp2* KO::PV-Cre mice injected with *Camkii*-GCAMP8m.
(C) Number of interaction bouts with the cage-mate or novel mouse during the social memory trial of the 4-Chamber social assay or WT::PV-Cre and *Mecp2* KO::PV-Cre mice injected with *Camkii*-GCAMP8m.
(D) Duration of interaction bouts with the cage-mate or novel mouse during the social memory trial of the 4-Chamber social interaction test for WT::PV-Cre and *Mecp2* KO::PV-Cre mice injected with *Camkii*-GCAMP8m.
(E) Number of interaction bouts during the sociability trial for WT::PV-Cre and *Mecp2* KO::PV-Cre mice injected with *FLEX*-GCAMP8f.
(F) Duration of interaction bouts during the sociability trial for WT::PV-Cre and *Mecp2* KO::PV-Cre mice injected with *FLEX*-GCAMP8f.
(G) Number of interaction bouts during the social memory trial for WT::PV-Cre and *Mecp2* KO::PV-Cre mice injected with *FLEX*-GCAMP8f.
(H) Duration of interaction bouts during the social memory trial for WT::PV-Cre and *Mecp2* KO::PV-Cre mice injected with *FLEX*-GCAMP8f.
(I) Velocity of WT::PV-Cre and *Mecp2* KO::PV-Cre mice during the social memory trial injected with *Camkii*-GCAMP8m (PYR expression) and *FLEX*-jRGECO1a (PV-IN expression)
(J) Distance traveled of WT::PV-Cre and *Mecp2* KO::PV-Cre mice during the social memory trial injected with*Camkii*-GCAMP8m (PYR expression) and *FLEX*-jRGECO1a (PV-IN expression). Data are mean ±SEM. *P<0.05, **P<0.01, ***P<0.001, ****P<0.0001.

**Supplemental Figure 3: Activity pattern analysis between WT::PV-Cre and *Mecp2* KO::PV-Cre using the AUC of jGCaMP8m transients to estimate the dynamics of mPFC PYR and PV-IN spiking activity**

(A) Simultaneous visualization of independent recordings of *Camkii*-GCaMP8m (PYR) activities from WT::PV-Cre and *Mecp2* KO::PV-Cre mice during behavioral interactions with the toy object in the sociability trial. Dark blue line denotes average signal from WT::PV-Cre mice, shaded blue denotes SEM. Dark red line denotes average signal from *Mecp2* KO::PV-Cre mice, shaded red denotes SEM. Time zero denotes behavioral bout onset

(B) mPFC PYR AUC analysis before entering the interaction zone (Pre, p=0.539) and following onset of investigation with the toy object (Post, p=0.506; n=8 *Mecp2* KO::PV-Cre vs. n=11 WT::PV-Cre)

(C) Simultaneous visualization of independent recordings of *Camkii*-GCaMP8m (PYR) activities from WT::PV-Cre and *Mecp2* KO::PV-Cre mice during behavioral interactions with the cage-mate in the sociability trial.

(D) AUC of mPFC PYR activities during interactions with the cage-mate (Pre, p=0.466; Post, p=0.698)

(E) Simultaneous visualization of independent recordings of *Camkii*-GCaMP8m (PYR) activities from WT::PV-Cre and *Mecp2* KO::PV-Cre mice during behavioral interactions with the cage-mate in the social memory trial.

(F) AUC of mPFC PYR activities during interactions with the cage-mate (Pre, p=0.122; Post, p=0.078; n=8 *Mecp2* KO::PV-Cre vs. n=11 WT::PV-Cre)

(G) Simultaneous visualization of independent recordings of *Camkii*-GCaMP8m (PYR) activities from WT::PV-Cre and *Mecp2* KO::PV-Cre mice during behavioral interactions with the novel mouse in the social memory trial.

(H) AUC of mPFC PYR activities during interactions with the novel mouse (Pre, p=0.017; Post, p=0.347)

(I) Simultaneous visualization of independent recordings of *FLEX*-GCaMP8f (PV-IN) activities from WT::PV-Cre and *Mecp2* KO::PV-Cre mice during behavioral interactions with the toy object in the sociability trial.

(J) AUC of mPFC PV-IN activities during interactions with the toy object (Pre, p=0.349; Post, p=0.111; n= 8 per group).

(K) Simultaneous visualization of independent recordings of *FLEX*-GCaMP8f (PV-IN) activities from WT::PV-Cre and *Mecp2* KO::PV-Cre mice during behavioral interactions with the cage-mate in the sociability trial.

(L) AUC of mPFC PV-IN activities during interactions with the cage-mate (Pre, p=0.323; Post, p=0.775)

(M) Simultaneous visualization of independent recordings of *FLEX*-GCaMP8f (PV-IN) activities from WT::PV-Cre and *Mecp2* KO::PV-Cre mice during behavioral interactions with the cage-mate in the social memory trial.

(N) AUC of mPFC PV-IN activities during interactions with the cage-mate (Pre, p=0.256; Post, p=0.457; n=8 per group).

(O) Simultaneous visualization of independent recordings of *FLEX*-GCaMP8f (PV-IN) activities from WT::PV-Cre and *Mecp2* KO::PV-Cre mice during behavioral interactions with the novel mouse in the social memory trial.

(P) AUC of mPFC PV-IN activities during interactions with the cage-mate (Pre, $p=0.268$; Post, $p=0.873$). Data are mean ± SEM. *$P<0.05$, **$P<0.01$, ***$P<0.001$, ****$P<0.0001$.

**Supplemental Figure 4: Activity pattern analysis between mPFC PYR and PV-IN population dynamics using the AUC of jGCaMP8m transients to estimate spiking transients in WT::PV-Cre and *Mecp2* KO::PV-Cre mice**

(A) Simultaneous visualization of independent recordings of *Camkii*-GCaMP8m (PYR) and *FLEX*-jGCaMP8f (PV-IN) in WT::PV-Cre mice calcium traces averaged across animals during behavioral interactions with the toy mouse in the sociability trial. Dark purple line denotes average signal from PYRs (*Camkii*), shaded purple denotes SEM. Dark green line denotes average signal from PV-INs (*PV*), shaded green denotes SEM. Time zero denotes behavioral bout onset

(B,C) AUC of mPFC PYR (*Camkii*) and PV-IN (*PV*) activities from WT::PV-Cre mice during interactions with the toy object (Pre, p=0.557;Post, p= 0.660; n=8-11 per group)

(D) Simultaneous visualization of independent recordings of *Camkii*-GCaMP8m (PYR) and *FLEX*-GCaMP8f (PV-IN) activities from WT::PV-Cre mice during behavioral interactions with the cage-mate in the sociability trial

(E,F) AUC of mPFC activities in WT::PV-Cre mice (Pre, p=0.161; Post, p=0.237) during interactions with the cage-mate in the sociability trial

(G) Simultaneous visualization of independent recordings of *Camkii*-GCaMP8m (PYR) and *FLEX*-GCaMP8f (PV-IN) activities from WT::PV-Cre mice during behavioral interactions with the cage-mate in the social memory trial

(H,I) AUC of mPFC activities in WT::PV-Cre mice (Pre, p=0.322; Post, p=0.132; n=8-11 per group) during interactions with the cage-mate in the social memory trial

(J) Simultaneous visualization of independent recordings of *Camkii*-GCaMP8m (PYR) and *FLEX*-GCaMP8f (PV-IN) activities from WT::PV-Cre mice during behavioral interactions with the novel mouse in the social memory trial

(K,L) AUC of mPFC activities in WT::PV-Cre mice (Pre, p=0.077; Post, p=0.906) during interactions with the novel mouse in the social memory trial

(M) Simultaneous visualization of independent recordings of *Camkii*-GCaMP8m (PYR) and *FLEX*-GCaMP8f (PV-IN) activities from *Mecp2* KO::PV-Cre mice during behavioral interactions with the toy in the sociability trial

(N,O) AUC of mPFC activities in *Mecp2* KO::PV-Cre mice during interactions with the toy object (Pre, p=0.317.; Post, p=0.367; n= 8-11 per group)

(P) Simultaneous visualization of independent recordings of *Camkii*-GCaMP8m (PYR) and *FLEX*-GCaMP8f (PV-IN) activities from *Mecp2* KO::PV-Cre mice during behavioral interactions with the cage-mate in the sociability trial

(Q,R) AUC of mPFC activities in *Mecp2* KO::PV-Cre mice during interactions with the cage-mate (Pre, p=0.255; Post, p=0.487)

(S) Simultaneous visualization of independent recordings of *Camkii*-GCaMP8m (PYR) and *FLEX*-GCaMP8f (PV-IN) activities from *Mecp2* KO::PV-Cre mice during behavioral interactions with the cage-mate in the social memory trial

(T,U) AUC of mPFC activities in *Mecp2* KO::PV-Cre mice during interactions with the cage-mate (Pre, p=0.0019; Post, p=0.0067; n=8-11 per group).

(V) Simultaneous visualization of independent recordings of *Camkii*-GCaMP8m (PYR) and *FLEX*-GCaMP8f (PV-IN) activities from *Mecp2* KO::PV-Cre mice during behavioral interactions with the novel mouse in the social memory trial

(W,X) AUC of mPFC activities in *Mecp2* KO::PV-Cre mice during interactions with the cage-mate (Pre, p=0.5141; Post, p=0.175). Data are mean ± SEM. *P<0.05, **P<0.01, ***P<0.001, ****P<0.0001

**Supplemental Figure 5 (Related to Figures 5,6)**

(A) Representative histological image of GCaMP8f and jRGECO1a and fiber placement for dual-color fiber photometry in the mPFC. Scale bar, 100 µm.

(B) Time spent interacting with the toy mouse or cage-mate during the sociability trial of the 4-Chamber social interaction test for WT::PV-Cre mice

(C) Time spent interacting with the the toy mouse or cage-mate during the sociability trial of the 4-Chamber social interaction test for *Mecp2* KO::PV-Cre mice

(D) Discrimination indices of the sociability trial of *Mecp2* KO::PV-Cre and WT::PV-Cre mice

(E) Time spent interacting with the cage-mate or novel mouse during the social memory trial of the 4-Chamber social interaction test for WT::PV-Cre mice

(F) Time spent interacting with the the cage-mate or novel mouse during the social memory trial of the 4-Chamber social interaction test for *Mecp2* KO::PV-Cre mice

(G) Discrimination indices of the social memory trial of *Mecp2* KO::PV-Cre and WT::PV-Cre mice
