## Supplemental Figures for "Altered activity of mPFC pyramidal neurons and parvalbumin-expressing interneurons during social interactions in a *Mecp2* mouse model for Rett syndrome"

### Supplemental Figure 1

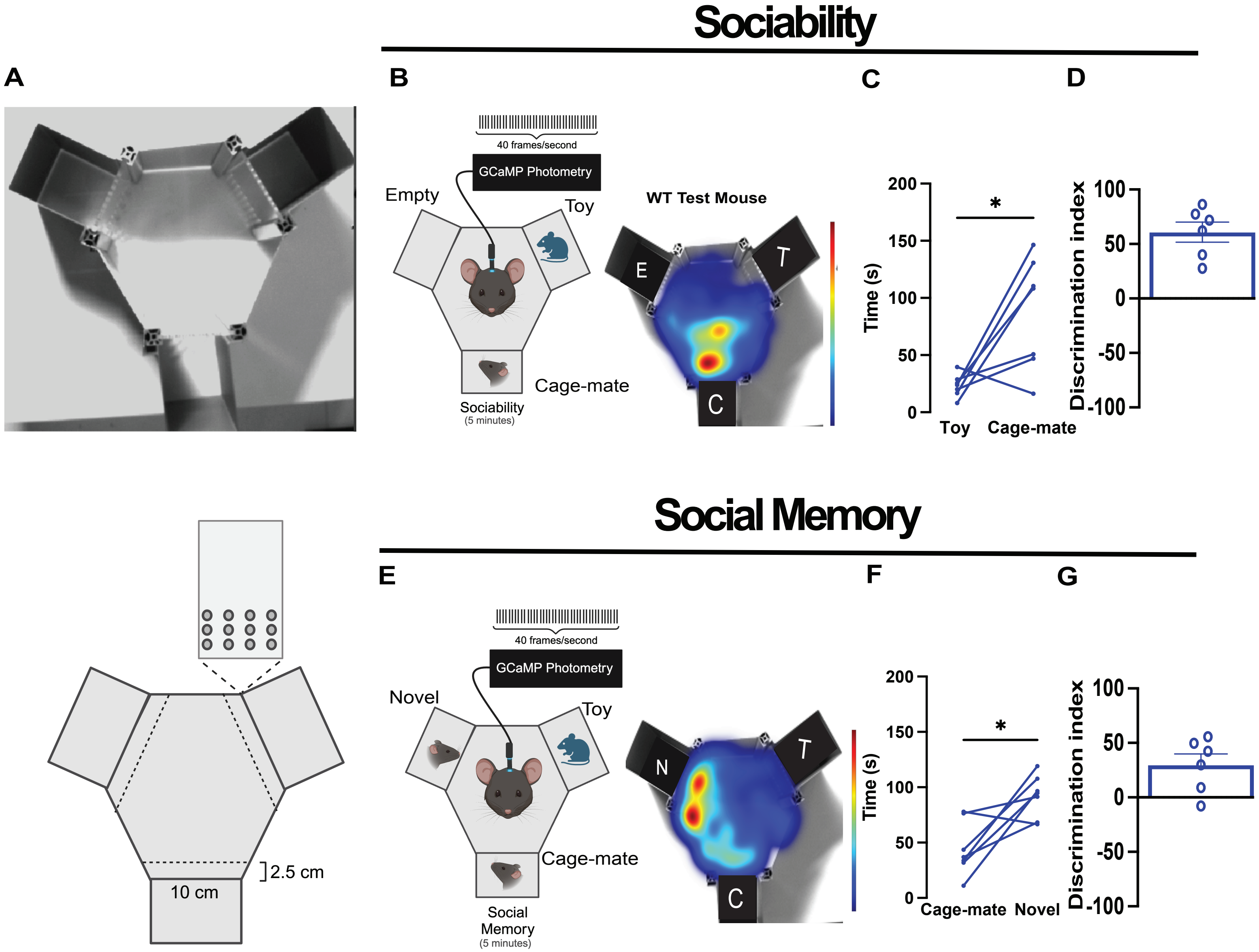

### Supplemental Figure 2

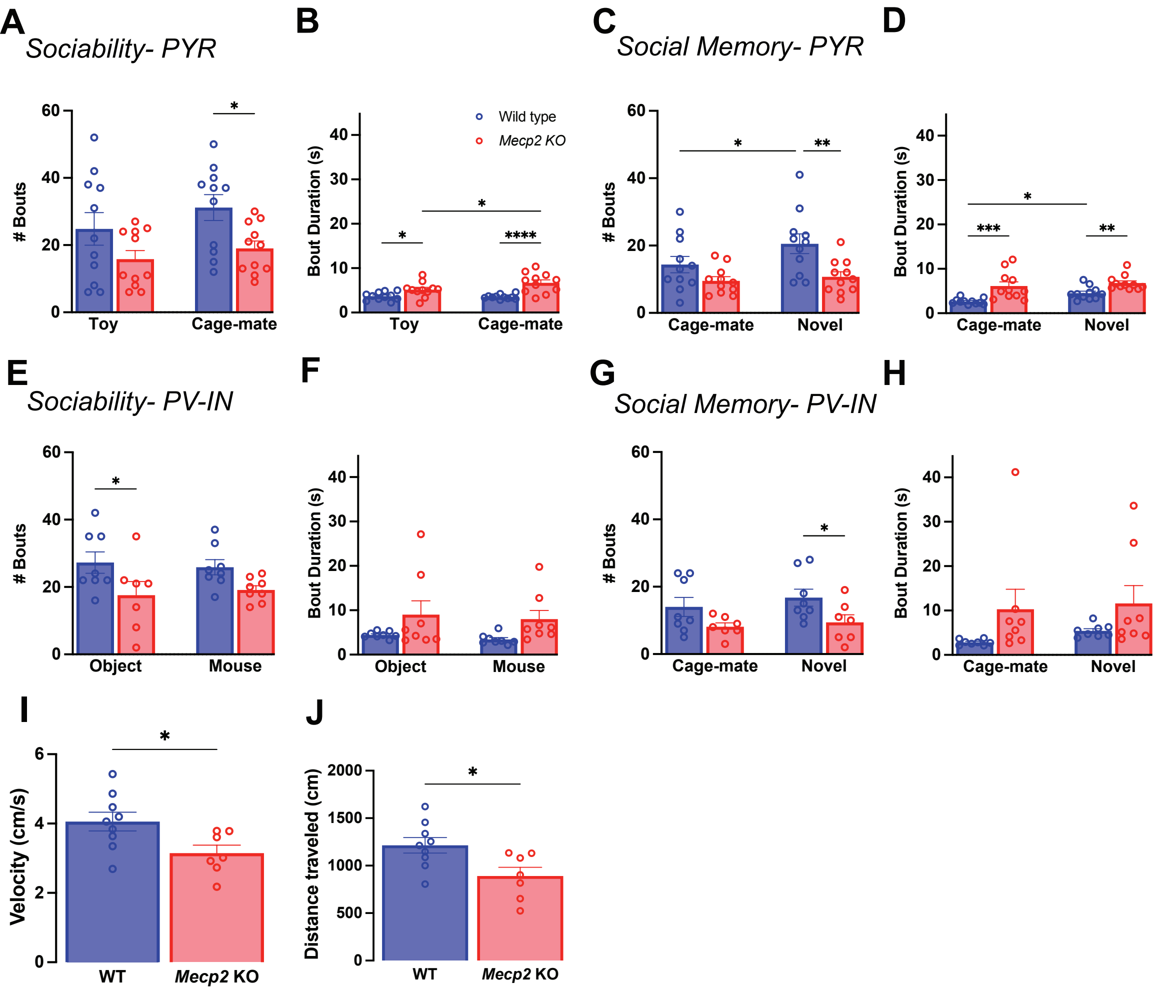

**Supplemental Figure 3** AUC Sociability PYR- WT vs. KO

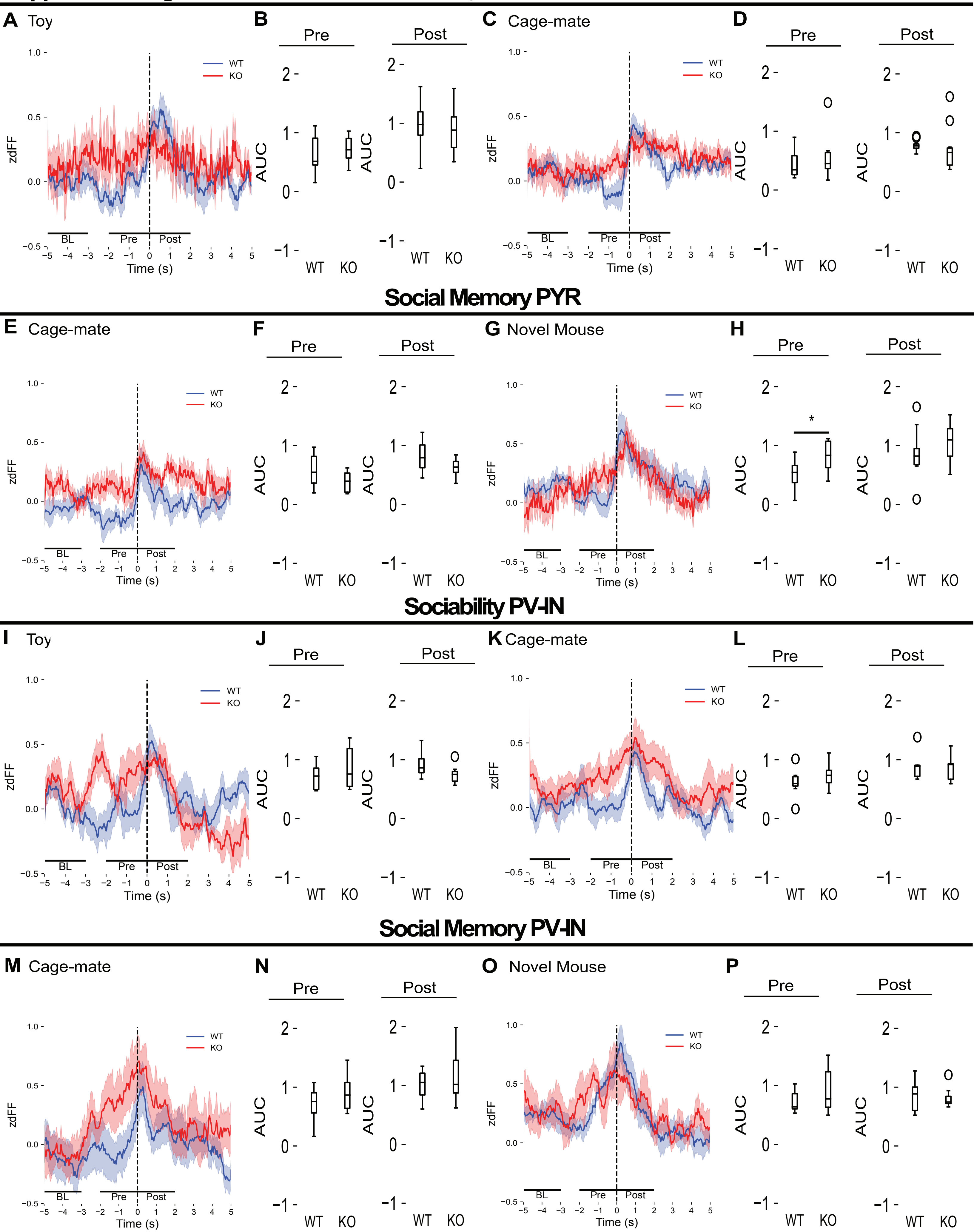

**Supplemental Figure 4**

#### AUC Sociability WT PYR vs. PV-IN

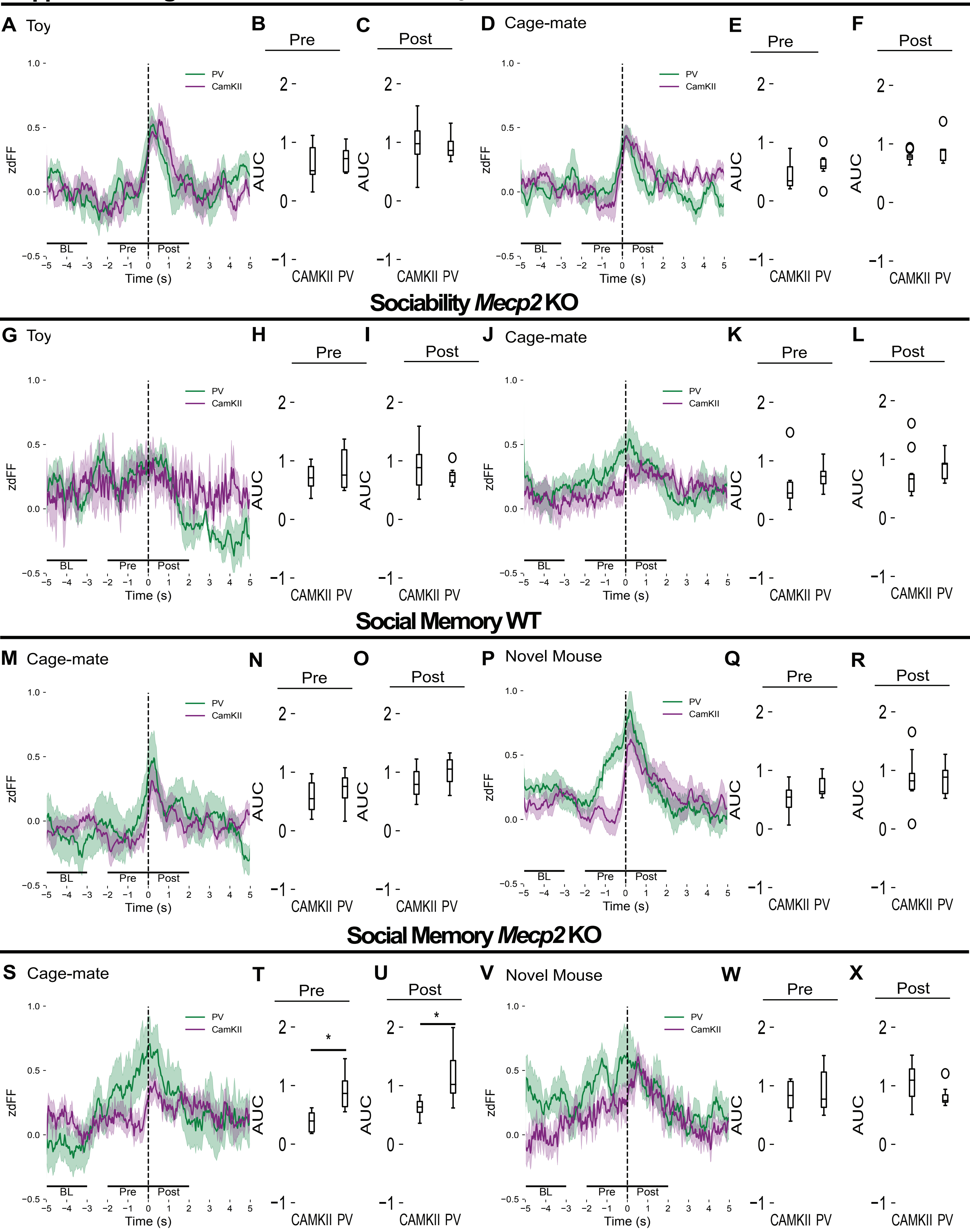

### Supplemental Figure 5 (Related to Figures 5 and 6)

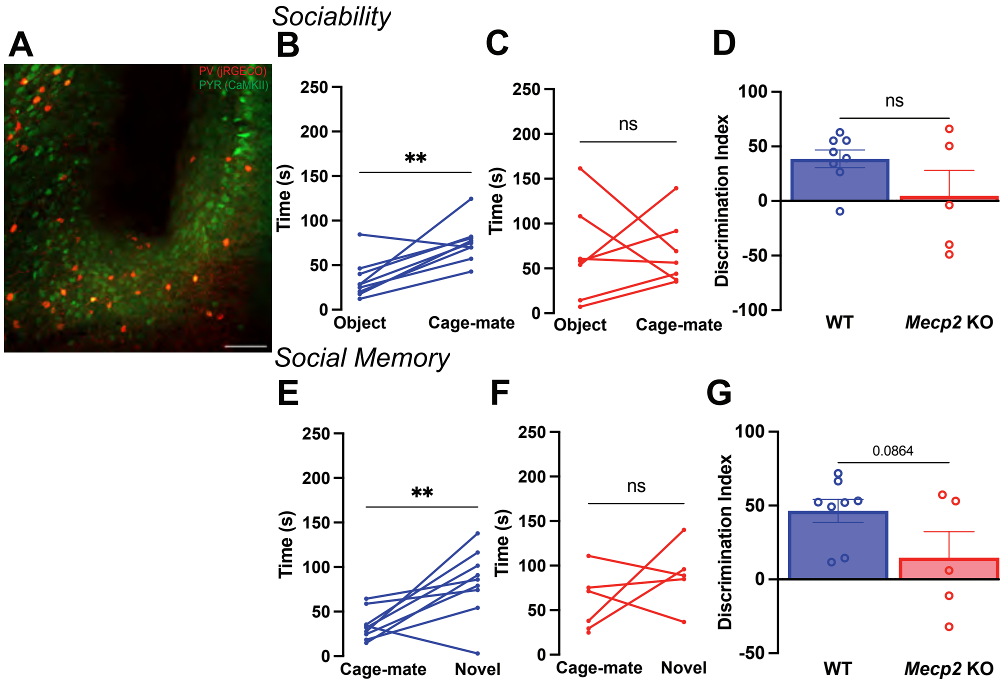
